# Variations in Atlantic water influx and sea-ice cover drive taxonomic and functional shifts in Arctic marine bacterial communities

**DOI:** 10.1101/2022.08.12.503524

**Authors:** Taylor Priest, Wilken-Jon von Appen, Ellen Oldenburg, Ovidiu Popa, Sinhué Torres-Valdés, Christina Bienhold, Katja Metfies, Bernhard M. Fuchs, Rudolf Amann, Antje Boetius, Matthias Wietz

## Abstract

The Arctic Ocean is experiencing unprecedented changes as a result of climate warming, necessitating detailed analyses on the ecology and dynamics of biological communities to understand current and future ecosystem shifts. Here we show the pronounced impact that variations in Atlantic water influx and sea-ice cover have on bacterial communities in the East Greenland Current (Fram Strait) using two, 2-year high-resolution amplicon datasets and an annual cycle of long-read metagenomes. Densely ice-covered polar waters harboured a temporally stable, resident microbiome. In contrast, low-ice cover and Atlantic water influx shifted community dominance to seasonally fluctuating populations enriched in genes for phytoplankton-derived organic matter degradation. We identified signature populations associated with distinct oceanographic conditions and predicted their ecological niches. Our study indicates progressing “Biological Atlantification” in the Arctic Ocean, where the niche space of Arctic bacterial populations will diminish, while communities that taxonomically and functionally resemble those in temperate oceans will become more widespread.

## INTRODUCTION

The Arctic Ocean is experiencing unprecedented changes as a result of climate warming. Of particular significance is the rapid decline in sea-ice extent and thickness^1,2^, with future projections indicating a potential for frequent ice-free summers by 2050^3^. In the Eurasian Arctic, accelerated rates of sea-ice decline have been linked to an increase in heat transport from inflowing Atlantic waters^4^, which together weaken water column stratification and increase vertical mixing. The expanding influence of Atlantic water in the Arctic Ocean, termed Atlantification, not only impacts hydrographic conditions but also provides avenues for habitat range expansion of temperate organisms^5,6^. The consequences of such perturbations on Arctic Ocean ecology are expected to be considerable. In order to predict and understand future changes in the ecosystem state and functioning of the Arctic Ocean, research on the ecology and dynamics of biological communities at the interface between Arctic and temperate oceans is essential^7^.

In seasonally ice-covered areas of the Arctic Ocean, bacterial communities exhibit seasonal temporal dynamics, similar to those in temperate ecosystems. These patterns are driven by pulses of organic matter released from phytoplankton blooms^8^ and the melting of first year sea-ice^9^. In recent decades, declining sea-ice cover has extended the growing season and increased open-water habitat space of phytoplankton, resulting in a 30% increase in net annual primary production between 1998 and 2012^10^ in shelf and slope areas of the Arctic. Phytoplankton bloom phenology is also shifting, with secondary autumn blooms now being observed in seasonally ice-covered areas^11^. These changes will alter the production and availability of organic matter to bacterial communities over spatial and temporal scales. Recent evidence has shown that sea-ice dynamics also influence the availability of organic matter in surface waters and the transport of carbon and microorganisms to the deep-sea^12–14^. Generally, ice margins are highly productive, because of the combination of early light availability, stratification and diatom-based blooms, which produce large particles and result in relatively high carbon export ^15^. However, strong melt events can intensify stratification, resulting in low nutrient supply and delayed export. For example, such a phenomenon in the Fram Strait was found to slow the biological carbon pump by up to 4 months compared to an ice-free situation^12^. Furthermore, declining sea-ice and warming Atlantic waters favour smaller flagellates, resulting in a pelagic retention system with reduced total annual export^16^. Thus, understanding how sea-ice dynamics influence bacterial communities will provide insights into future biological and biogeochemical changes.

The Fram Strait, the main deep-water gateway between the Arctic and Atlantic Oceans, is a key location for conducting long-term ecological research over environmental gradients and under changing conditions^17^. Fram Strait harbours two major current systems; the East Greenland Current (EGC), which transports polar water southwards in its upper layer, and the West Spitsbergen Current (WSC) that transports Atlantic water northward. The EGC accounts for the export of ∼50% of freshwater and ∼90% of sea-ice from the central Arctic Ocean and carries Arctic hydrographic signatures^18^. Large-scale recirculation of Atlantic water (AW) into the EGC by eddies is a continuously occurring process, although the magnitude varies across latitudes and over temporal scales^19,20^. The mixing of these water masses in the marginal ice zone (MIZ) creates different hydrographic regimes reflective of Arctic, mixed water and Atlantic conditions, which can harbour unique bacterial compositions^21,22^. Carter-Gates *et al*.^21^ predicted that future Atlantification of the Arctic may result in a shift towards temperate, Atlantic-type communities. However, further assessments of microbial population dynamics across different temporal and spatial scales, i.e. under Arctic vs Atlantic conditions, are needed to validate such hypotheses.

Here, we performed a high-resolution analysis of the temporal variation of bacterial taxonomy and function at two locations in the EGC between 2016-2018 (MIZ) and 2018-2020 (core EGC), covering the full spectrum of ice cover, daylight and hydrographic conditions (Arctic to Atlantic water masses). Our study was enabled by the “Frontiers in Arctic Marine Monitoring” (FRAM) Ocean Observing System that employs mooring-attached sensors and autonomous Remote Access Samplers (RAS) to continuously monitor physiochemical processes and biological communities at the LTER HAUSGARTEN in the Fram Strait. This analysis encompasses a 2 two-year 16S rRNA amplicon dataset supplemented with an annual cycle of long PacBio HiFi read metagenomes, expanding a previous assessment of microbial dynamics over a single annual cycle in the EGC^23^. We hypothesise that high AW influx and low sea-ice cover will result in bacterial communities dominated by chemoheterotrophic populations, resembling those of temperate ecosystems. Our investigation provides essential insights into the effects of the changing Arctic on marine microbial ecology and biogeochemical cycles.

## RESULTS AND DISCUSSION

The amplicon dataset incorporates samples (>0.2 µm fraction) collected at weekly to biweekly intervals at the Marginal Ice Zone (MIZ; 2016–2018) and in the core of the EGC (core-EGC; 2018– 2020), between 70–90 m depth (Supplementary Table S1). The two locations were selected in order to capture the full spectrum of water mass and sea ice conditions. The core-EGC featured dense ice cover (in the following abbreviated with “high-ice”) and predominantly polar water (PW) year-round. In contrast, the MIZ featured variable, generally lower ice cover (in the following abbreviated with “low-ice”) and Atlantic water (AW) influx (Figure 1). To provide a more visual representation, animated GIFS were created for current velocities at the depth of sampling (Supplementary Figure S1) and sea-ice cover dynamics (Supplementary Figure S2) over the four year period. By combining the high-resolution data from both mooring locations, we are able to assess bacterial community dynamics over temporal scales and in relation to contrasting Arctic- and Atlantic-dominated conditions.

**Figure 1.**
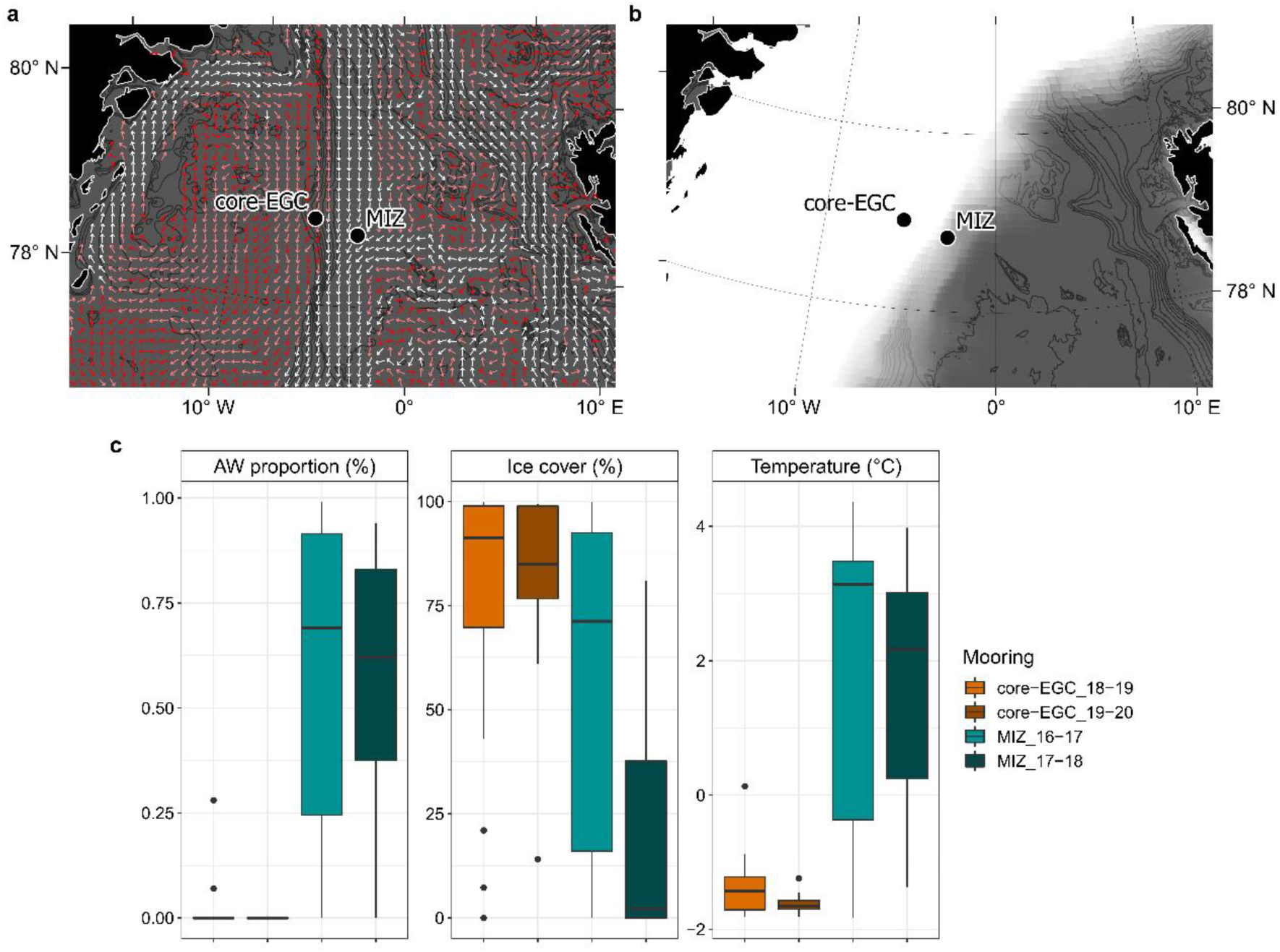
Geographical location of seafloor moorings and variation in environmental conditions in MIZ (2016–2018) and core-EGC (2018–2020). **a)** Example representation of monthly average (January 2020) current velocities at the approximate depth of water sampling (78m). White and dark red arrows indicate strongest and weakest velocities, respectively. **b)** Example representation (December 2019) of sea-ice cover. Increasing opacity of white colour reflects increasing sea-ice cover, where pure white = 100% sea-ice cover. Values for current velocities and sea-ice concentration were obtained from copernicus.eu under the ‘ARCTIC_ANALYSIS_FORECAST_PHY_002_001_a’. **c)** Boxplots illustrating variation in AW proportion, ice cover and temperature at the moorings. The bathymetric map was made using publically available bathymetry data from GEBCO [17].

### Bacterial community and population dynamics over temporal scales

Combining the 2 two-year amplicon datasets resulted in 3988 non-singleton Amplicon Sequence Variants (ASVs) (Supplementary Table S2) being recovered, which were initially used in a taxonomy-independent approach to assess community dynamics over environmental gradients. Shifts in bacterial community composition with changing environmental conditions were evident (Figure 2). A stepwise significance test identified AW proportion, daylight and past ice cover (average ice cover of the seven days preceding the sampling event) as the significant factors explaining compositional variation (model R^2^ = 0.23, p-value = 0.001) (Supplementary Table S1). AW proportion explained 13% of the total variation, compared to 6% for daylight and 4% for past ice cover. The pronounced impact of AW reflects previous observations that different water masses harbour distinct bacterial assemblages^21,25,26^ and has important implications for the future Arctic Ocean, as AW influence is expected to expand. The impact of daylight can be directly and indirectly linked to bacterial community dynamics, through their own phototrophic capacities and indirectly via phytoplankton dynamics. Here, changes in light availability could reflect seasonal dynamics, which are well evidenced in temperate and polar ecosystems^27–29^ and also occur in ice-free areas of the Fram Strait^30^. The influence of ice cover on light availability highlights the importance of sea-ice cover for shaping bacterial communities in the Arctic.

**Figure 2.**
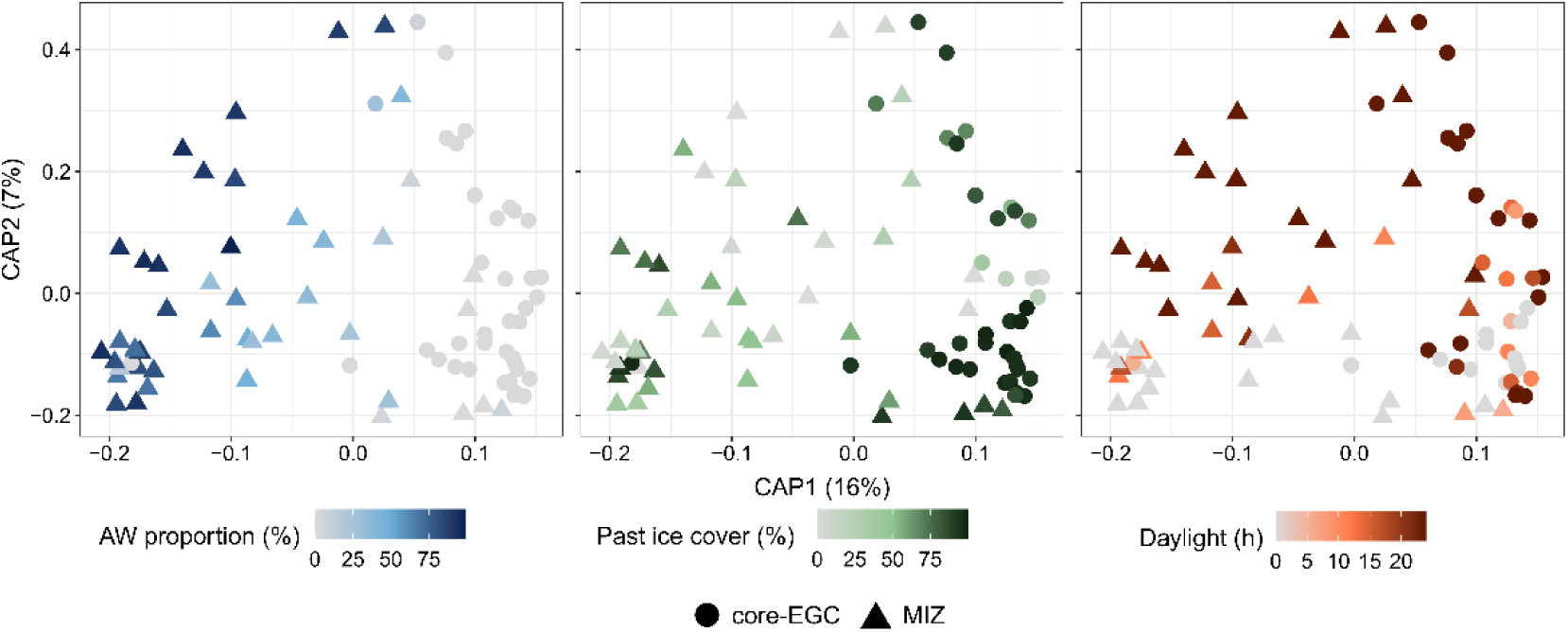
Community structure across variations in water mass, ice cover and daylight conditions. Distance-based redundancy analysis based on Bray-Curtis dissimilarities of community composition along with AW proportion (blue symbols), past ice cover (green symbols) and daylight (orange symbols) as constraining factors. The factors were selected using a stepwise significance test and combined into a single model (R^2^ = 0.1, pvalue = 0.01) that constrains 14% of the total variation. For ease of interpretation, the environmental conditions are visualised individually on the same ordination.

To gain a more detailed understanding on bacterial community structuring in the EGC, the dynamics of ASVs across samples was assessed. In total, 75% of the ASVs were detected at both mooring sites, whilst 16% and 8% were unique to the MIZ and core-EGC, respectively. The frequency of detection and maximum relative abundance of ASVs shared by both sites exhibited a strong positive linear relationship, i.e. those identified in more samples also reached higher maximum relative abundances (Figure 3a). To facilitate further comparisons, we categorised ASVs into three groups: a) resident ASVs (Res-ASVs), present in >90% of samples, b) intermittent ASVs (Int-ASVs), present in 25 - 90% of samples, and c) transient ASVs (Trans-ASVs), present in <25% of samples. There were only 232 Res-ASVs, but these represented the largest proportion of the sampled bacterial communities (48 - 88% relative abundance). In comparison, the 1904 Int-ASVs constituted 12 – 48% and the 1852 Tran-ASVs between 0.4 – 5.8% of relative abundances. Presence of a dominant resident microbiome, represented by a minority of ASVs, corresponds to previous observations in summertime Fram Strait samples^31^ as well as the Western English Channel and Hawaiian Ocean time-series^27,32^.

**Figure 3.**
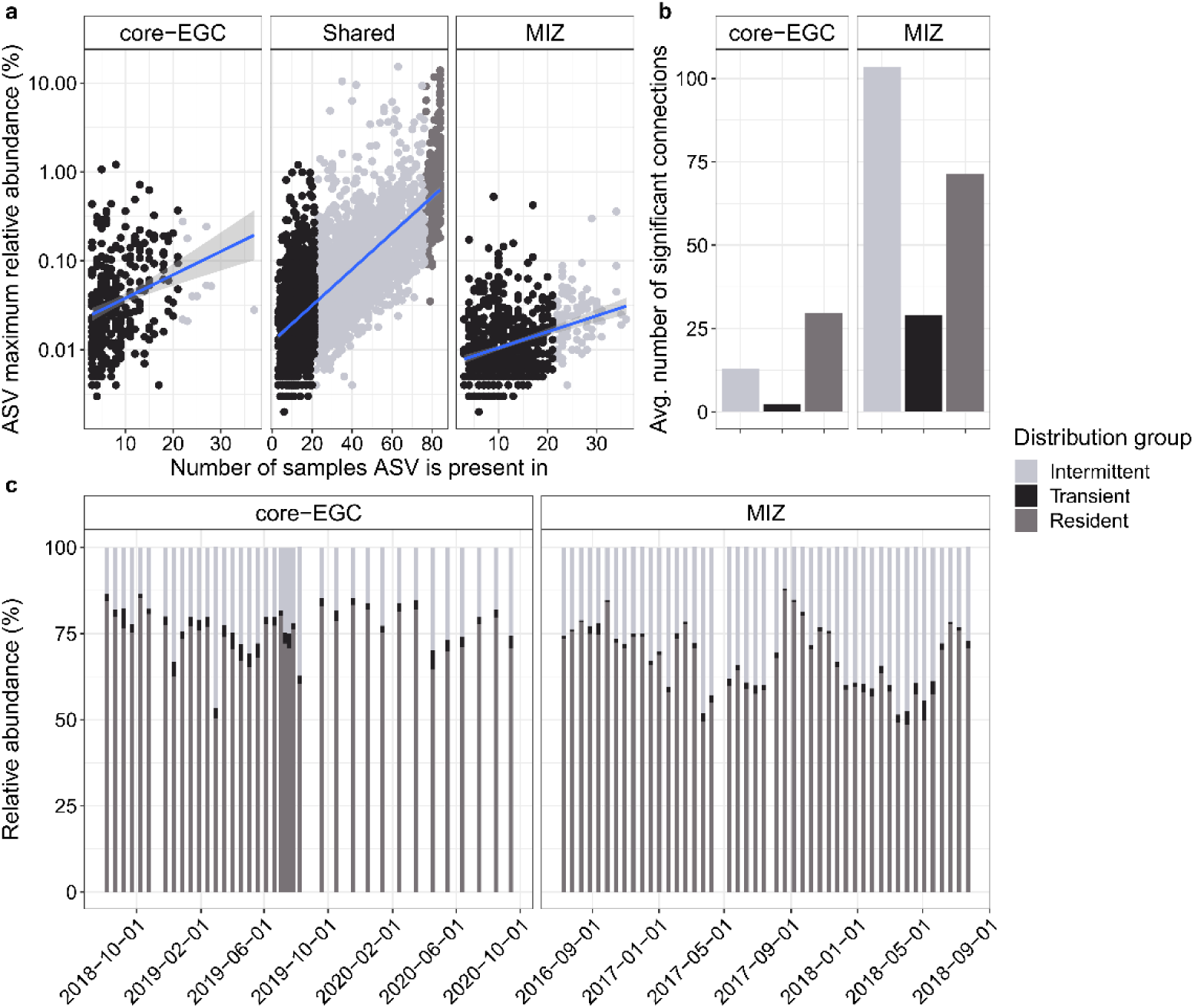
Distribution dynamics and co-occurrence of ASVs. **a)** The occurrence of ASVs across samples in relation to their maximum relative abundances along with categorization into resident, intermittent and transient. **b)** Average number of connections within the co-occurrence networks for resident, intermittent and transient ASVs. **c)** Relative abundance dynamics of the resident, intermittent and transient ASVs over time.

The fluctuations in abundance of the three community fractions were primarily associated with changes in AW proportion, with strong negative correlations for the resident (Pearson’s coefficient: −0.50, p-value <0.01) and transient (Pearson’s coefficient: −0.32, p-value < 0.01) microbiomes and strong positive correlations for the intermittent fraction (Pearson’s coefficient 0.57, p-value <0.01). Visualising the dynamics of the community fractions further highlights this association, with the resident microbiome exhibiting a more stable pattern in core-EGC compared to MIZ samples (Figure 3c). The resident microbiome was phylogenetically diverse and incorporated both abundant and rare community members. Res-ASVs were assigned to 47 families and 79 genera, with the *Flavobacteriaceae* (n=15), *Magnetospiraceae* (n=13), *Marinimicrobia* (n=11) and SAR11 Clade I (n=22) and Clade II (n=17) harbouring the largest diversity. Maximum relative abundances of Res-ASVs ranged from 0.035 – 13.9%, with the most prominent being affiliated with the SAR11 Clade Ia (asv1 - 14%), *Polaribacter* (asv6 - 14%), *Aurantivirga* (asv7 - 12%), SUP05 Clade (asv2 - 12%), SAR92 Clade (asv16 - 11%) and SAR86 Clade (asv3 – 9%). Pronounced fluctuations of the intermittent community (11 – 48% relative abundance) coincided with AW influx at the MIZ. The intermittent community was more phylogenetically diverse than the resident microbiome, encompassing 250 genera, and also comprised rare and abundant populations that reached between 0.004 – 15% maximum relative abundance. The most diverse taxa included the SAR11 Clade II (n=148), *Marinimicrobia* (n=129), NS9 Marine Group (n=78), AEGEAN-169 (n=73) and *Nitrospinaceae* (n=56). Those reaching the largest relative abundances were affiliated with *Luteolibacter* (asv24 - 15%), *Flavobacterium* (asv140 - 10%), *Polaribacter* (asv206 - 10%) and *Colwellia* (asv89 - 9%). The resident and intermittent community fractions shared 71 genera, which constitutes 90% of the genus-level diversity of the resident microbiome. Hence compositional changes over temporal scales are driven by dynamics on the species- and population-level.

To consolidate the observed community structuring and further illuminate ASV dynamics, we computed co-occurrence networks and contextualised them with environmental conditions (Supplementary Figure S3). Networks were computed for both moorings separately in order to assess patterns in microbial population dynamics under Arctic- and Atlantic-dominated conditions. In the core-EGC network, Res-ASVs exhibited twofold more significant co-occurrences than Int-ASVs, averaging 29 and 13 respectively (Figure 3b). In contrast, Int-ASVs were more connected in the MIZ network. This pattern further illustrates the stability of the resident microbiome under Arctic-dominated conditions. In both networks, a distinct cluster of co-occurring, summerly-peaking (June-August) ASVs were observed. These included ASVs reaching the highest relative abundances, such as asv24-*Luteolibacter* (Int-ASV), asv6-*Polaribacter* (Res-ASV) and asv7-*Aurantivirga* (Res-ASV). The MIZ network exhibited additional strong seasonal structuring. No further co-occurrence patterns in relation to environmental conditions were observed in the core-EGC network (Supplementary Figure S3).

Overall, the seasonally and spatially variable conditions in the MIZ drive substantial dynamics of distinct bacterial populations. Accordingly, the environmentally less dynamic core-EGC is reflected in a more stable resident microbiome that fluctuates less with seasons, likely reflecting adaptations to polar water and nearly year-round ice cover.

#### Taxonomic signatures of distinct environmental conditions

A sparse partial least squares regression analysis (sPLS) identified 430 ASVs that were associated with distinct environmental conditions. These ASVs formed eight distinct clusters based on similar, significant correlations to seven environmental parameters (thresholds: coefficients >0.4, p-values <0.05) (Figure 4a). The composition of ASVs in each cluster were largely unique, revealing distinct taxonomic signatures of specific environmental conditions (Figure 4b). The three largest clusters incorporated 88% of the ASVs and were separated based on their associations to different water mass and ice cover conditions. Clusters C1 and C2 represented AW conditions, with C1 also being associated with low-ice cover. In contrast, cluster C8 represented polar water (PW), with high-ice cover. In accordance with the distribution dynamics described above, the AW-associated clusters comprised a higher proportion of Int-ASVs, 51 – 88%, compared to ∼50% Res-ASVs in PW-associated clusters. An additional five smaller clusters (C3 - C7) were also identified that corresponded to polar day and polar night under different ice cover and water mass conditions. Comparing the most prominent ASVs of each cluster (reaching >1% relative abundance) revealed unique taxonomic signatures at the genus-level (Figure 4b). For instance, *Amylibacter*, SUP05 Clade and AEGEAN-169 were signatures of the AW-associated, low-ice cluster C1, whilst the SAR324 Clade, NS2b Marine Group and *Magnetospira* were signatures of the PW-associated, high-ice cluster C8. Furthermore, the PW-associated cluster C8 harboured a distinctly larger phylogenetic diversity compared to the other clusters. Overall, this pattern underlines water mass and ice cover as major drivers of community structure, whilst a smaller number of ASVs are strongly influenced by daylight and seasonality.

**Figure 4.**
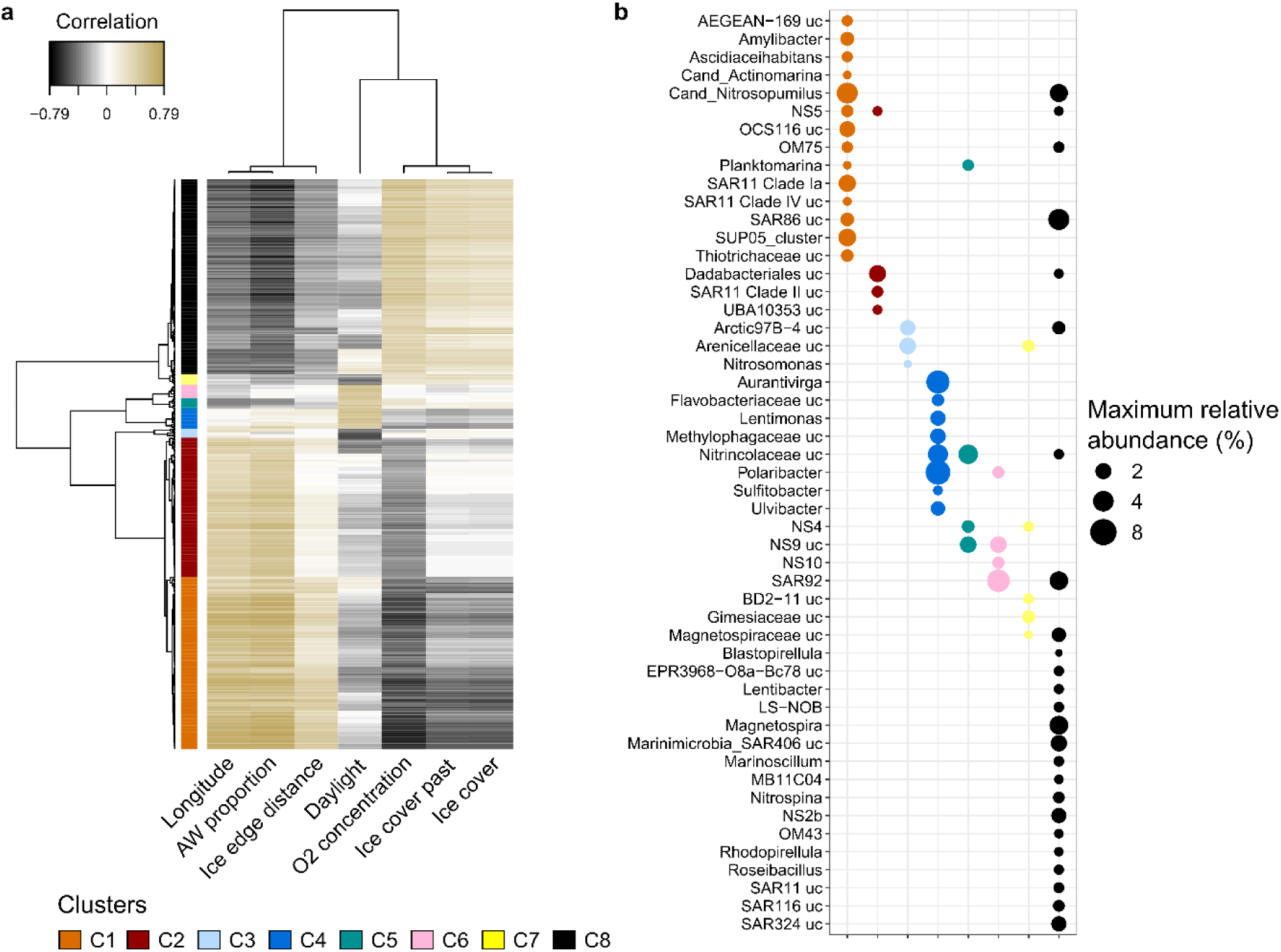
Sparse partial least square regression linking community structure and environmental parameters. **a)** Heatmap showing the eight major sPLS clusters identified that encompass 430 ASVs with significant correlations to environmental conditions. **b)** Representation of the most prominent genera in each cluster. ASVs that did not reach >1% were excluded from the taxonomic comparison, whilst the remaining were grouped by genus and the maximum abundance of each genus shown. Due to high collinearity with AW proportion, temperature and salinity were excluded from sPLS analysis.

### Signature populations associated with distinct environmental conditions

We contextualized ASV dynamics with metagenome-assembled genomes (MAGs) to link distribution with metabolic potential and subsequently predict ecological niches of populations under different environmental conditions. From nine PacBio HiFi read metagenomes, derived from the 2016-17 annual cycle in the MIZ, we recovered 43 manually-refined, population-representative MAGs (delineating cut-off of 98% average nucleotide identity; ANI). The MAGs represented 26 – 49% of the metagenomic reads. Of these, twelve were high-quality drafts whilst the remainder were medium-quality drafts^33^. Despite this, medium-quality MAGs were highly contiguous (average number of contigs = 33) and >80% contained at least one complete rRNA gene operon. The generated MAGs represented a broad phylogenetic diversity, including 35 genera, 27 families and nine classes (Figure 5 and Supplementary Table S3). Comparing the MAGs to those recently recovered from the Fram Strait^22^ indicated that 32 were novel species (<95% ANI), whilst eleven represented previously recovered populations (>99% ANI).

**Figure 5.**
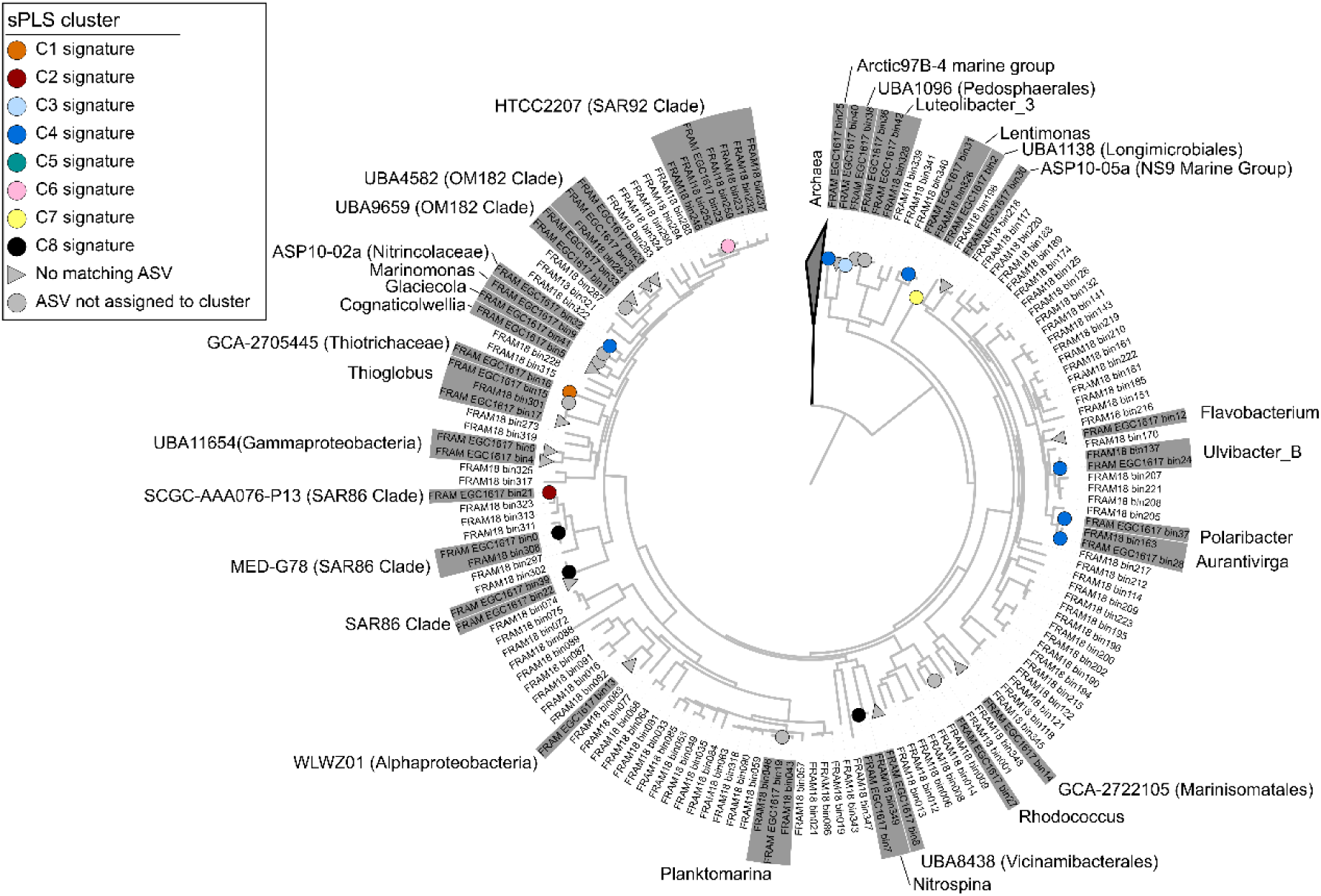
Ribosomal protein-based phylogenetic tree of MAGs from this study and those previously recovered from the Fram Strait. Tree is based on concatenated alignment of 16 ribosomal proteins, as previously used by Hug et al. [81]. MAGs from this study (indicated by colours circles) were combined with those previously published by Priest et al. [15]. Only MAGs that contained at least 8 ribosomal proteins were included in the tree. Grey boxes around labels indicate distinct genera with MAG representatives recovered in this study.

Through competitive read recruitment, 27 of the MAGs were linked to distinct ASVs (based on 100% identity threshold). Of these, 18 could be associated with sPLS clusters and thus distinct environmental conditions. These 18 representatives of sPLS clusters, which encompass an ASV and MAG, are hereon referred to as “signature populations” (Table 1 and Figure 5). Signature populations included some of the most prominent representatives of each cluster, such as asv6-*Polaribacter* and asv7-*Aurantivirga* from cluster C4 and asv18-SAR86 Clade from cluster C8. Based on their dynamics being driven primarily by ice cover and water mass, we define signature populations of cluster C1 and C2 as Atlantic signatures and those of C8 as Arctic signatures. For consistency, signature populations will be identified by their asv number, with the corresponding MAG name provided in Table 1.

**Table 1.**
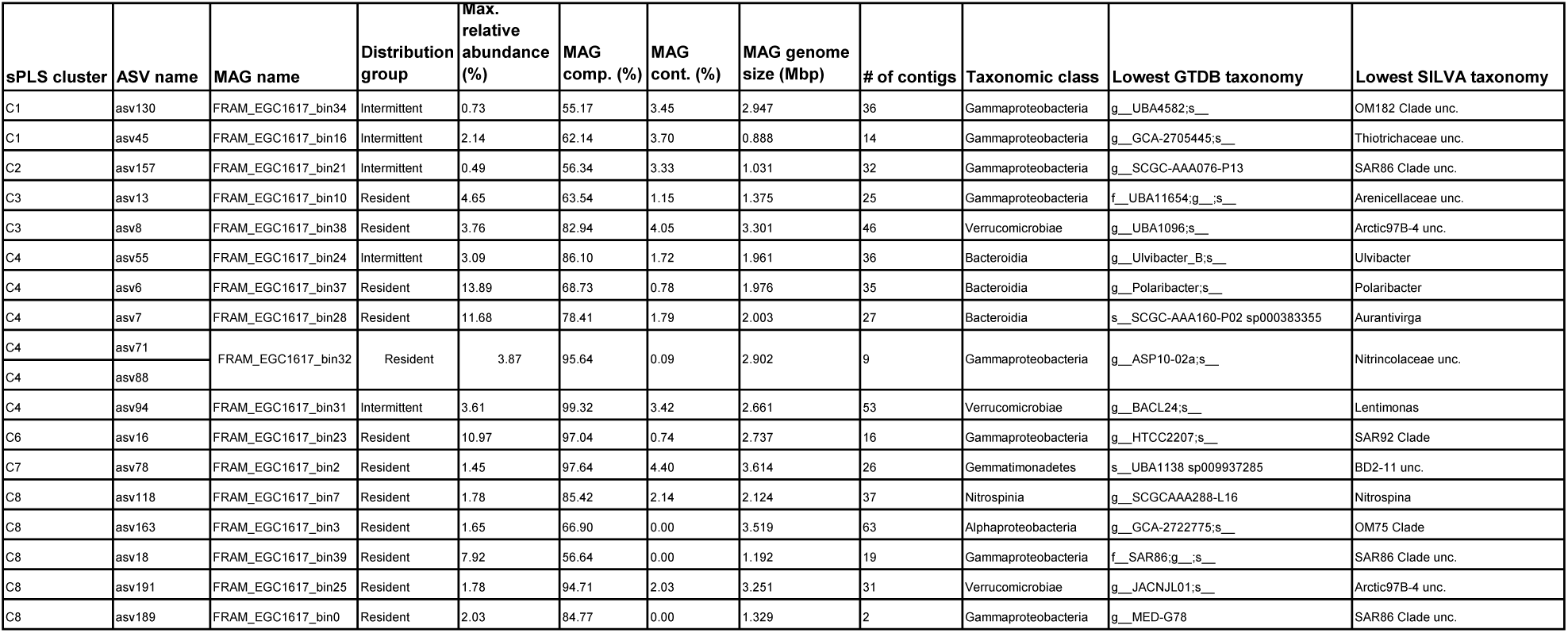
Taxonomic, genomic and distribution information on signature populations. Completeness (MAG comp.) and contamination (MAG cont.) were estimated using CheckM v1.1.3. MAGs were taxonomically classified using the GTDB r202 database with the GTDBtk tool v1.7.0. ASV taxonomy was assigned within the DADA2 pipeline using the SILVA 138 SSU Ref NR99 database.

### Distribution of signature populations across the Arctic Ocean

In order to validate our observations on signature population dynamics with environmental conditions and their assignment as resident, intermittent and transient, we assessed their distribution across the Arctic Ocean using the Tara Oceans prokaryote size fraction dataset (Supplementary Figure S4). Signature population MAGs of the polar day clusters (C4 and C6) were, on average, present at the highest relative abundances in upper euphotic zone samples (5-100 m depth). In particular, asv6-*Polaribacter* in cluster C4, with an average relative abundance of 2.7%, and asv16-SAR92 Clade in cluster C6, with an average relative abundance of 2.1%. At lower depths (>100 m), polar day signatures decreased and Arctic and polar night signature populations increased, i.e. as in C8 and C3, respectively. The most prominent were asv13-*Arenicellaceae* in cluster C3 and asv18-SAR86 Clade in cluster C8, with average relative abundances of 0.16 and 0.66% respectively. The Arctic Ocean sampling campaign of the Tara Oceans was conducted largely during summer months (May - October) and was restricted to locations above the continental shelf, which typically experience ice-free conditions or low-ice cover during that time period. As such, the higher prevalence of polar day signature populations (C4 and C6) in the upper water column is in agreement with their observed dynamics in the EGC. Water column stratification following sea-ice melt and polar day conditions likely restricts Arctic and polar night signature populations to deeper waters. These populations could be expected to increase in the upper water column in conjunction with deeper vertical mixing in the winter. Furthermore, it is likely that the “true” Arctic signature populations identified in our dataset are more prevalent in the Arctic Ocean basin, and are transported southward with water exiting the central Arctic through the EGC.

### Ecological niches of signature populations

By assessing temporal dynamics in functional genes, we are able to predict the ecological niches of signature populations within the context of the environmental conditions they represent. Of particular interest were the signature populations of Atlantic (clusters C1 and C2) and Arctic (cluster C8) conditions, as their ecology could provide insights into how bacterial community structure may shift under future conditions in the Arctic Ocean. Ecological descriptions of daylight driven clusters are provided in Supplementary Information S1 and tables with complete genome functional annotations for all signature populations are provided in Supplementary Files S1.

### Atlantic signature populations

Atlantic signature populations were affiliated with the *Gammaproteobacteria* class, and included representatives of the OM182 Clade (UBA4582), *Thiotrichaceae* (GCA-2705445) and the SAR86 Clade (SCGC-AAA076-P13). All three populations reached significantly higher relative abundance values in MIZ compared to core-EGC, however, within the MIZ, distinct dynamics were observed. The *Thiotrichaceae* (asv45) population peaked during the summer season (polar day) in AW conditions, whilst the SAR86 Clade (asv157) population peaked during the polar night in AW conditions. The dynamics of the OM182 Clade (asv130) population was related only to AW and not to daylight. Functional gene predictions revealed unique lifestyles and metabolic capacities for each population that provide insights into the key factors likely driving the observed dynamics. The focus of the following analyses is placed on the asv45 and asv130 populations, whilst insights into the function and ecology of the asv157 population is provided in Supplementary Information S1.

The *Thiotrichaceae* population harbours the capacity to utilise phytoplankton-derived organic compounds, the availability of which likely stimulates growth under polar day conditions. These include methanethiol and C1 compounds. Methanethiol originates from the demethylation of dimethylsulfoniopropionate (DMSP)^34^, an osmoprotectant produced by phytoplankton. DMSP concentrations in the Arctic Ocean show spatial variation and are influenced by water mass and sea-ice, with highest concentrations reported in areas directly influenced by Atlantic water inflow (western Eurasian Arctic)^35^. The concentration of DMSP in these regions is tightly coupled to chlorophyll *a*^35,36^. As such, methanethiol is likely more available in Atlantic waters under polar day conditions. The oxidation of methanethiol, through the methanethiol oxidase gene (MTO), results in the production of formaldehyde and hydrogen sulfide. In the asv45 population, we identified the MTO gene along with a complete tetrahydromethanopterin (H4-MPT)-dependent oxidation pathway and sulfide oxidation machinery (dsrAB and soeABC). Combined, this genetic repertoire would allow the asv45 population to use methanethiol as a carbon, sulfur and energy source.

A similar metabolism has been reported for members of the *Rhodobacteraceae*, which are capable of degrading methanethiol and subsequently oxidising the hydrogen sulfide for energy generation^37^. The H4-MPT-dependent formaldehyde oxidation pathway has been traditionally affiliated with methanogenic bacteria but was also demonstrated in methylo- and methanotrophic members of the Alpha- and *Gammaproteobacteria*^38^. To our knowledge, this is the first such description in a sulfur-oxidizing member of the *Thiotrichaceae* family. Although experimental evidence is needed to consolidate these findings, we obtained the species-representative genomes from the assigned GTDB genus (GCA-2705445), and confirmed the presence of the above-described pathways in each. The GCA-2705445 genus contains several representatives that are classified as *Thiothrix* in NCBI. The distinct metabolic features described above may represent unique characteristics and, in line with the GTDB classification, suggests that GCA-2705445 species are distinct from other *Thiothrix*.

The OM182 Clade population differed from the other AW-associated signatures by showing daylight-independent dynamics. Functional gene annotations indicated a motile lifestyle with the capacity to oxidise sulfur and carbon monoxide (CO) as well as degrade taurine and methylamine, thus representing an aerobic, sulfur-oxidising methylotroph. Furthermore, the presence of the complete *sox* system along with polysulfide reductase (*pshAB*) and flavocytochrome c-sulfide dehydrogenase (*fccAB*) genes indicates the capacity to store and use elemental sulfur. The diverse metabolic capacities of the asv130 population may explain the observed dynamics over the time-series, providing it the capacity to switch nutrient and energy sources under different conditions. For example, under high daylight conditions, CO oxidation combined with the utilisation of organic compounds presumably provides sufficient energy and nutrients for growth. CO production in the oceans is linked to the photolysis of coloured dissolved organic matter and direct production by phytoplankton^39,40^, and thus concentrations would be elevated during polar day and in periods of high productivity. Under such conditions, the capacity to use taurine and methylamine, which are compounds related to phytoplankton production and organic matter degradation respectively, would provide further access to carbon, nitrogen and sulfur. CO oxidation as a supplemental energy source has been previously evidenced in some marine *Gammaproteobacteria*^41^, however the dominant organisms performing such processes are typically affiliated with *Rhodobacteraceae* members. In general, only a few heterotrophic populations inhabiting the upper water column have been linked to sulfur- and CO-oxidation. The OM182 Clade may be an important contributor to the biogeochemical cycling of some carbon and sulfur species in the Arctic pelagic environment.

### Arctic signature populations

The Arctic signature populations each exhibited highly similar dynamics with peak relative abundances under high-ice cover, low AW proportion and low daylight conditions. The most prominent of these were the asv18 (SAR86 Clade), asv118 (*Nitrospina*) and asv163 (OM75 Clade) (Figure 6). All five populations harboured distinct metabolic capacities that were either indicative of chemoautotrophic lifestyles or chemoheterotrophic lifestyles, with a capacity to use diverse substrates for growth beyond phytoplankton-derived organic compounds. In this regard, their genetic tools were notably different to Atlantic signature populations. Here with cluster C8 signatures, we focussed on the SAR86 Clade population (due to the high relative abundance) and on the Arctic97B-4 populations (due to the limited ecological information currently available). Detailed descriptions of other signature populations are given in Supplementary Information S1. In addition, we assessed the cluster C7 signature population, asv78 – BD2-11, as it also represents PW under low daylight conditions and is affiliated with a rather unknown but resident taxon.

**Figure 6.**
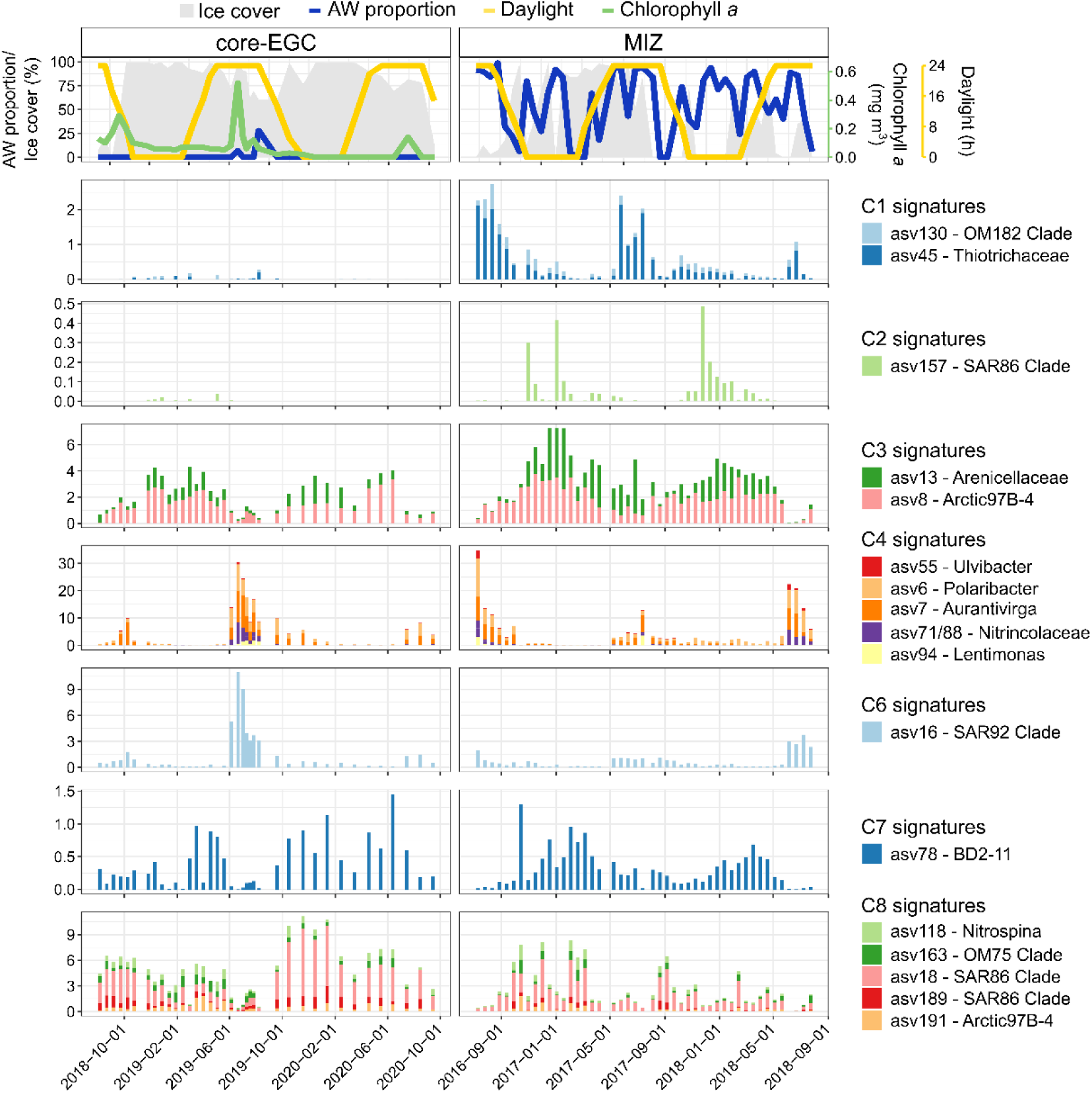
Temporal dynamics of signature populations. Signature populations were identified as ASV representatives from sPLS clusters that a corresponding MAG was recovered for (based on 100% identity threshold competitive read recruitment). The temporal dynamics visualized are derived from ASV data only. The missing chlorophyll *a* data in 2016-18 is due to the lack of a sensor on the MIZ mooring.

The most prominent Arctic signature population, asv18, likely represents a novel genus in the SAR86 Clade, based on <60% average amino acid identity (AAI) to other GTDB representatives. However, it shares >99% AAI to a recently recovered MAG from the Fram Strait (FRAM18_bin252)^22^, which corroborates its assignment to the resident microbiome. SAR86 Clade members are known as photoheterotrophs, with distinct ecotypes relating to phototrophic and carbohydrate degradation capacities^42^. They are also known as one of the most prominent gammaproteobacterial responders to spring phytoplankton blooms in temperate ecosystems, wherein they are predicted to use carbohydrates and DMSP^43^. In agreement, our Arctic MAG encodes a green-light proteorhodopsin, typical of lower photic zone-inhabiting organisms^44^, as well as carbohydrate degradation genes. However, the larger peptidase (n=19) to CAZyme (n=7) gene count ratio and the affiliation of CAZyme gene families with peptidoglycan recycling (GH103 and GH84), suggests a preference for proteinaceous substrates. Furthermore, the population encoded the capacity to metabolise D-amino acids, via conversion to α-keto acids^45^. In conjunction with the phylogenetic distance, these metabolic distinctions to other SAR86 Clade members may represent features of a novel, Arctic-specific genus.

The asv191 signature Arctic population was assigned to the Arctic97B-4 group (*Verrucomicrobiae*), for which no functional information is available to date. 16S rRNA gene-based studies have indicated elevated proportions of Arctic97B-4 in subsurface waters and a tight coupling with other deep water clades, such as SAR202^46,47^. An enrichment of Arctic97B-4- affiliated sequences was also identified in the small particle-attached fraction of Southern Ocean samples^48^. In accordance with these findings, the functional annotations of asv191 population suggest a motile chemomixotroph with the capacity to oxidise methane, fix carbon and degrade sulfated carbohydrates. Comparable to other marine *Verrucomicrobia*, the asv191 population encoded a high number of CAZymes (23 genes) and sulfatases (84 genes). However, the peaks of asv191 under no- to low-daylight conditions suggest that alternative substrates are also used. We identified the key marker genes for carbon fixation through the reductive TCA cycle (*korAB, por, trfAB*) and the aerobic oxidation of methane through formate and formaldehyde (*mdh, metF* and *folD*). However, the key methane monooxygenase gene (*pmo*) was not detected. This genetic repertoire is identical to a recently described MAG from the same family (*Pedosphaeraceae*), recovered from a bioreactor community^49^. The authors of that study suggested a potentially novel methane monooxygenase gene that was originally annotated as hypothetical. The aerobic oxidation of methane in the water column is typically associated with environments above continental shelves and oxygen minimum zones, where methane is supplied from the sediment or anaerobic processes below. Studies from above continental shelves in the Arctic have shown supersaturation of methane, with significantly elevated concentrations under sea-ice compared to ice-free conditions^50,51^. The increased prevalence of the Arctic97B-4 population under high-ice cover may thus be related to increased methane availability in conjunction with their particle-attached lifestyle, as methane production is known to occur in marine particulate organic matter^52^.

The cluster C7 representative, asv78, was assigned to two distinct classes, the BD2-11 terrestrial group (SILVA) and the *Gemmatimonadetes* (GTDB). This discrepancy reflects the relatively recent assignment and largely unresolved phylogeny of the *Gemmatimonadota* phylum. Based on the few available cultured representatives and 16S rRNA-gene based studies, the *Gemmatimonadetes* harbour aerobic/semi-aerobic chemoorganoheterotrophs inhabiting soil environments, but are also reported from freshwater habitats and deep-sea sediments^53,54^. Their presence in the upper marine water column however, is rarely reported. The asv78 population encodes the capacity to use a wide range of organic substrates for growth, and also perform aerobic denitrification. However, the presence of periplasmic nitrate reductase (*nap*) and nitrite reductase (*nirK*) genes but absence of downstream genes required for further reduction to N2 suggests an incomplete denitrification pathway. In addition, we identified genes to metabolise taurine, hypotaurine, D-amino acids, dicarboxylic acids and halogenated haloaliphatic compounds. The sources of these compounds in the marine environment vary, with taurine being attributed to phytoplankton and metazoa, D-amino acids to bacteria, and halogenated compounds to all forms of biota. With asv78’s capacity to reduce nitrate, this would provide metabolic flexibility to prevail under low daylight conditions and high-ice cover.

#### Whole community functional shifts with contrasting conditions

To find out if the functionality of the whole community also shifts with changes in ice cover and AW influx, we assessed differences in functional gene composition at the community level using the raw PacBio HiFi reads from the 2016 - 2017 MIZ samples. To date, only one study has published such metagenomic data from Arctic seawater samples^22^. From the nine metagenomes generated here, 17.6 million open reading frames were identified (Supplementary Table S4), of which 54% were assigned a function and 92% were assigned a taxonomy. Expectedly, taxonomic classifications of individual genes, varied in resolution, with 92% being assigned to a kingdom, 64% to a family and 37% to a genus. However, the GTDB-based pipeline employed here provided a wealth of taxonomic information that could not be obtained using the NCBI-nr database, an approach typically employed by published tools (Supplementary Information S1). The robustness of classifications was consolidated by comparing community composition recovered from reads and the ASVs, which showed high congruence at the class level (Supplementary Figure S5).

A dissimilarity analysis of whole-community functionality separated samples into two distinct clusters largely corresponding to high and low-ice cover (Figure 7). Permutation ANOVA revealed ice cover as a statistically significant factor (F-statistic = 12.6, p-value=0.009). A differential abundance analysis on normalized gene counts identified 1088 differentially abundant genes between these two clusters, with 328 and 845 genes enriched under high- and low-ice conditions, respectively. Gene enrichment was related to substrate uptake and degradation (Figure 8). Bacterial communities under low-ice exhibited an enhanced capacity to utilize carbohydrates, dissolved organic nitrogen (DON) and sulfur (DOS) compounds. Genes for nitrate utilisation were enriched under high-ice compared to ammonium utilisation under low-ice (Figure 9). High-ice conditions were enriched in genes involved in the metabolism of amino acids, proteins, aromatics and ketone compounds. These patterns reflect fundamental differences in community functionality, corroborating MAG-derived evidence that low-ice communities likely rely on labile organic compounds derived from phytoplankton.

**Figure 7.**
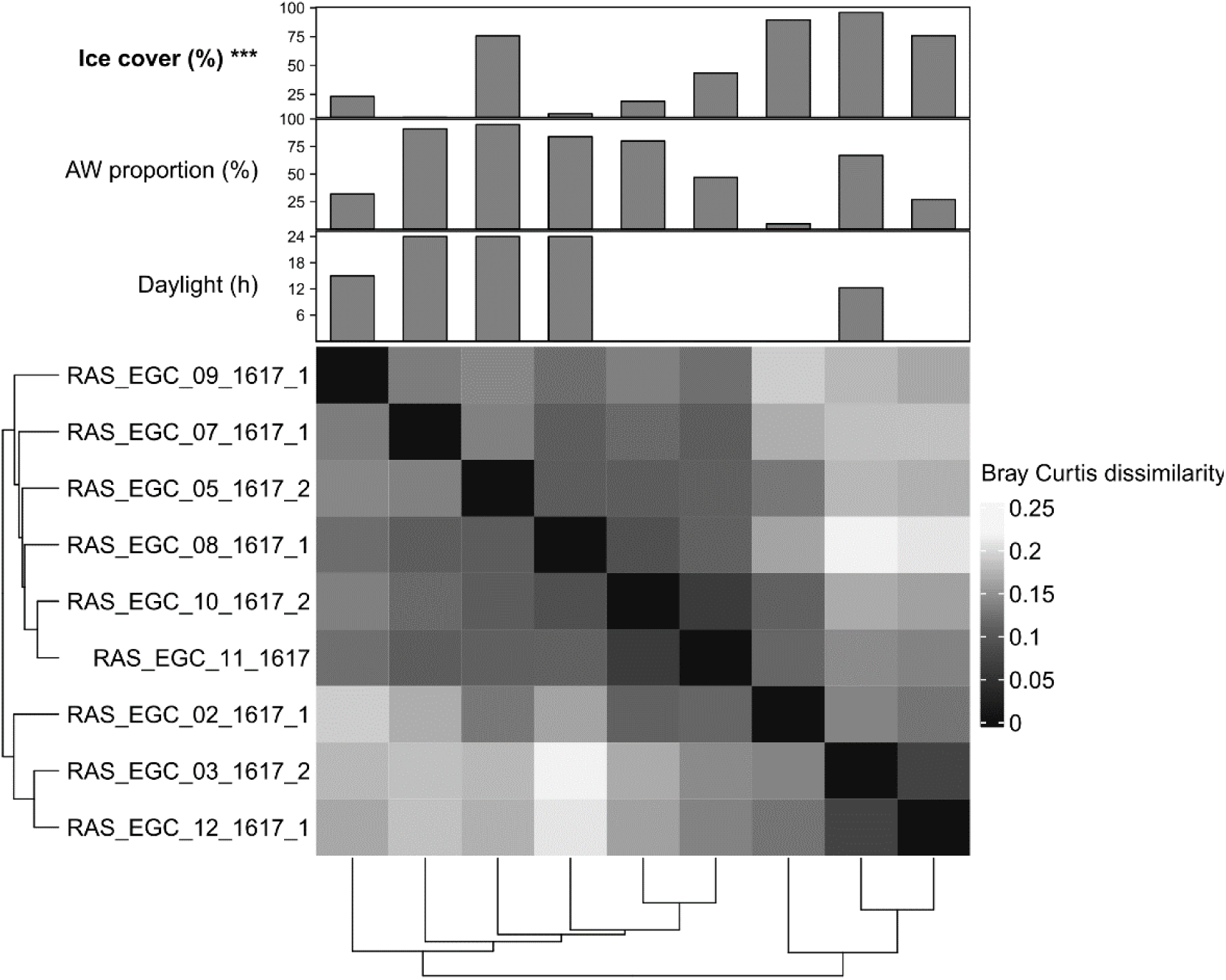
Clustering of sample functional gene composition based on Bray-Curtis dissimilarities.

**Figure 8.**
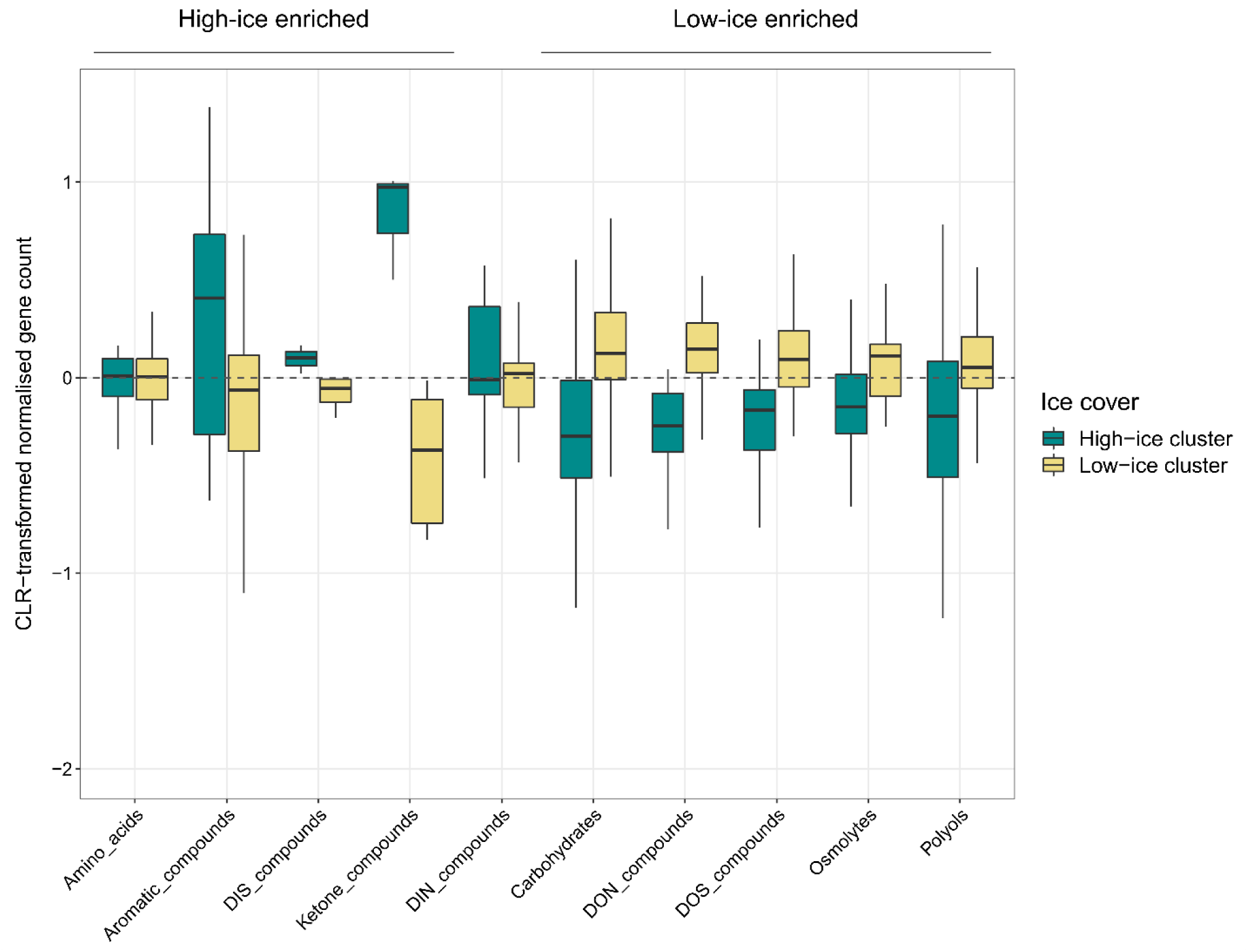
Summary of substrate uptake and degradation-related genes enriched with high- and low-ice coverage. Enriched genes were individually searched in Biocyc and KEGG, and subsequently placed into broader categories based on the type of substrate metabolism they were involved in. Values shown are centered-log ratio transformed normalised gene counts, as used in the differential abundance analysis.

**Figure 9.**
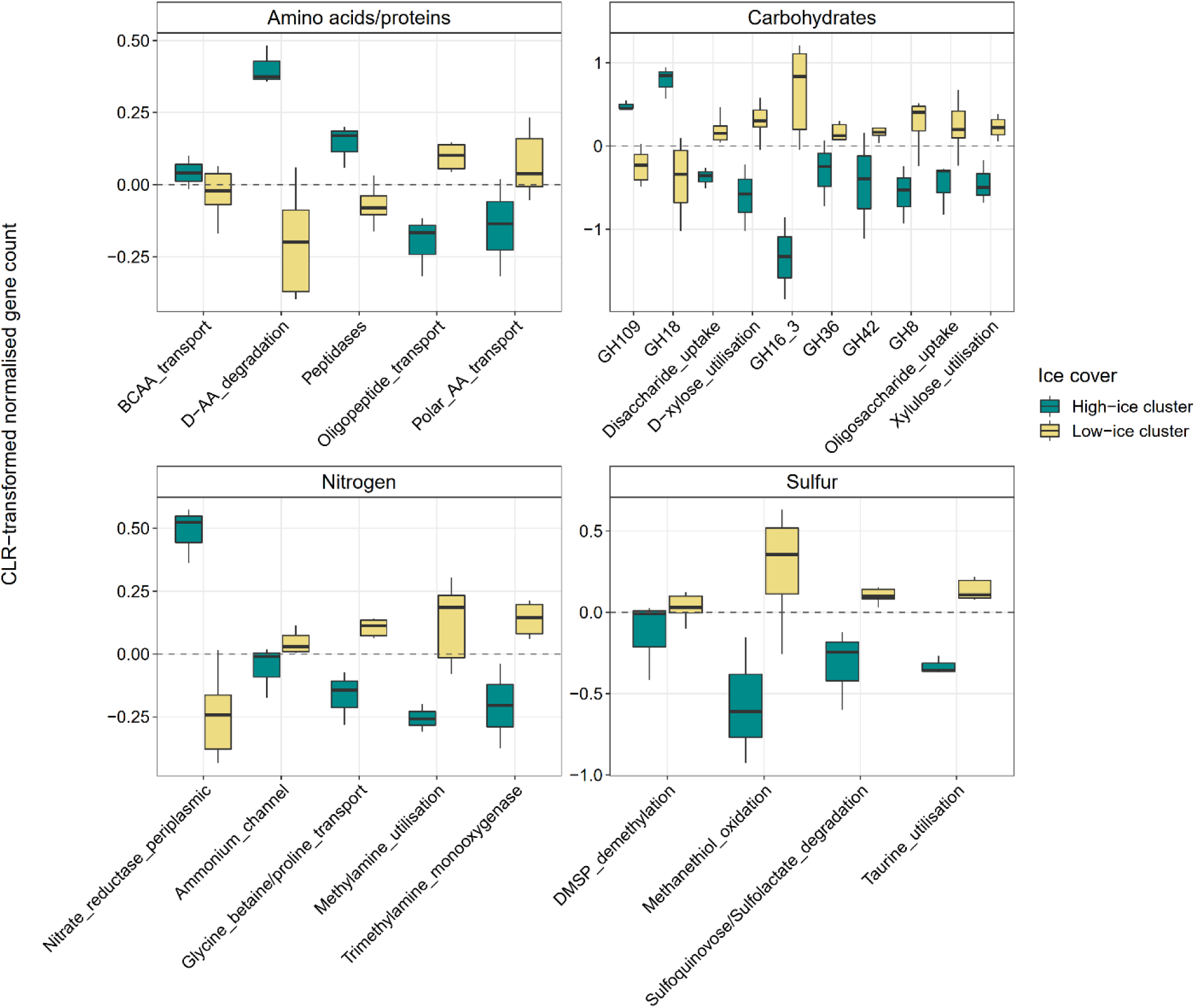
Selected genes involved in the uptake and degradation of organic and inorganic compounds enriched under high- and low-ice conditions. Enriched functional genes are displayed as the centered-log ratio transformed normalized gene counts. Where several genes of a single pathway or mechanism were identified as enriched, they were grouped into one and the term ‘utilisation’ used (e.g. “taurine_utilisation” indicates the uptake and degradation of taurine). When single genes were identified, the corresponding gene names are included.

### Functions enriched under low-ice cover

Enrichment of glycoside hydrolase families GH16_3, GH36, GH42 and GH8 indicate an increased potential to degrade laminarin and α-galactose- and β-galactose-containing polysaccharides (Figure 9). In addition, numerous transporters and degradation genes related to mono- and disaccharides were enriched, including for D-xylose, glucose and rhamnose. Carbohydrates represent a major component of dissolved organic matter, 15 - 50%^55,56^, and particulate organic matter, 3 - 18%^57^, and are key substrates for heterotrophic marine bacteria. The major source of carbohydrates in the oceans is phytoplankton, wherein they can constitute up to 50% of the cell biomass^58^. The release of carbohydrates during phytoplankton blooms stimulates the growth of heterotrophic bacteria, resulting in deterministic and recurrent dynamics driven by their carbohydrate utilisation capacity^29,59^. Phytoplankton production is also the primary source of other organic compounds, particularly DOS, such as DMSP, taurine and sulfoquinovose^60^. Although many bacteria harbour the capacity for inorganic sulfate assimilation, this process is energetically expensive. As such, using DOS compounds reduces energetic requirements and can additionally provide access to other nutrients, such as taurine that can act as a carbon, nitrogen and sulfur source. Additional genes, either directly or indirectly related to phytoplankton production and degradation, were also enriched, such as methylamine.

### Functions enriched under high-ice cover

50% fewer genes were enriched under dense ice, and they were mostly restricted to the recycling of bacterial cell wall carbohydrates, proteins, amino acids, aromatics and ketone compounds. Under high-ice cover, phytoplankton are less prominent^30^, limiting the availability of fresh labile organic matter and necessitating alternative growth strategies. For instance, the enrichment of a nitrate reductase gene indicates a specific potential to use inorganic substrates (Figure 9).

Enrichment of GH families 109 and 18, related to peptidoglycan degradation, suggests recycling of bacterial cell wall components as carbon and energy sources. GH18 is also known to contain chitinases^61^ and thus could indicate the degradation of chitin-rich materials such as carapaces and fecal pellets^62^. An increased reliance on bacterial-derived organic matter is further supported by the enrichment of D-amino acid degradation-related genes (Figure 9), as D-enantiomers of amino acids are largely derived from bacteria^63^. The enrichment in peptidases indicates that proteinaceous compounds play a more prominent role under high-ice, likely related to the production of related substrates by almost all organisms and hence wider availability.

Furthermore, we observed enrichment in genes for the degradation of aromatic and ketone compounds. Aromatic compounds in the Arctic Ocean typically originate from terrestrial organic matter that is sourced from rivers, constituting up to 33% of all Arctic Ocean DOM^64^. 12 – 41% of this terrestrial-derived DOM is exported to the North Atlantic via the EGC^64^. Therefore, the enrichment of genes for aromatic compound degradation indicates adaptations towards more diverse substrates under high-ice cover. These observations match reports for the *Chloroflexi* (SAR202) phylum inhabiting Arctic surface waters^65^, which were enriched in aromatic compound degradation genes compared to their deep-water counterparts. Together, these features suggest the presence of a community that “recycles” available substrates and is not reliant on fresh labile-organic matter from phytoplankton.

### Shifts in eukaryotic communities further support changes in ecosystem functioning

The observed ecosystem shifts between high and low-ice cover was reflected in the composition of eukaryotes at the same nine time points. High-ice cover conditions harbored increased proportions of *Dinophyceae, Syndiniales* and RAD-C radiolarians (Supplementary Figure S6), corresponding to prior reports of autumn-winter Arctic eukaryotic communities being dominated by heterotrophic-mixotrophic taxa ^30,66^. Although the proportions of diatoms (Bacillariophyta) were comparable at high- and low-ice, clear distinctions occurred at higher taxonomic resolution. High-ice conditions coincided with increased proportions of ice-associated taxa such as *Bacillaria, Naviculales* and *Polarella*. In contrast, the open-water diatoms *Thalassiosira* and *Pseudo-nitzschia* prevailed under low-ice cover, resembling temperate Atlantic phytoplankton communities. Also *Phaeocystis* was enriched under AW conditions. Overall, the predominance of ice-associated algae and heterotrophic-mixotrophic eukaryotes under dense ice, compared to pelagic diatoms under low-ice cover is also reflecting the difference in ecosystem functioning in different ice and daylight regimes.

## CONCLUSION

Climate change is amplified in the Arctic Ocean region, driving fundamental shifts in oceanographic and biological regimes. Here we show that variations in sea-ice extent and influx of Atlantic water masses considerably affect bacterial community dynamics and functionality. Densely ice-covered polar waters harbour a temporally stable, resident microbiome, which “recycles” available substrates and contains enriched signatures of autotrophic and inorganic substrate metabolism. In contrast, the ice margin with low-ice cover and more Atlantic water is dominated by seasonally fluctuating chemoheterotrophic populations, many of which are functionally linked to phytoplankton-derived organic matter. Both at population and community level, sea-ice cover had the strongest influence on bacterial functionality. Hence, we predict that the future Atlantification of the Arctic Ocean and continued reduction in sea-ice cover will shrink the ecological niches of signature Arctic populations, likely restricting them to the central basin and the core of the East Greenland current where ice is transported into the Atlantic. These shifts represent a “Biological Atlantification” in the Arctic Ocean that will have implications on future ecosystem functioning and carbon cycling.

## METHODS

### Seawater collection and processing

Autonomous sample collection and subsequent processing of samples proceeded according to Wietz et al.^23^. Briefly, Remote Access Samplers (RAS; McLane) were deployed over four consecutive annual cycles between 2016-2020, with deployments and recovery occurring each summer (recover of 2019/2020 mooring occurred in 2021). From 2016 – 2018, RAS were deployed in the MIZ (78.83° N −2.79° E) and from 2018 – 2020 in the core EGC (79°N −5.4°E) at nominal depths of 80 and 70m, respectively. Sampling occurred at weekly to biweekly intervals (Supplementary Table S1). At each sampling event, ∼1 L of seawater was autonomously pumped into sterile plastic bags and fixed with mercuric chloride (0.01% final concentration). After RAS recovery, water was filtered onto 0.22 µm Sterivex cartridges directly frozen at −20 °C until DNA extraction.

### Amplicon sequencing and analysis

DNA was extracted using the DNeasy PowerWater kit (Qiagen, Germany), followed by amplification of 16S rRNA gene fragments using primers 515F–926R^67^. Sequencing was performed on a MiSeq platform (Illumina, CA, USA) using 2 × 300 bp paired-end libraries according to the “16S Metagenomic Sequencing Library Preparation protocol” (Illumina). Amplicons were subsequently processed into amplicon sequence variants using DADA2. Analysis of ASV dynamics and subsequent generation of plots was performed in RStudio^68^, using primarily, the vegan^69^, limma^70^, mixOmics^71^, ggplot2^72^ and ComplexHeatmap^73^ packages. Briefly, community composition was compared using Bray-Curtis dissimilarities and distance-based redundancy analysis with the functions *decostand* and *dbrda* in vegan and visualised using ggplot2. The influence of environmental variables on community dissimilarity was determined using the *ordiR2step* and *anova*.*cca* functions in vegan. ASV dynamics across the time-series and assignment into distribution groups, e.g. resident, was determined by extracting information from the relative abundance matrix produced by DADA2. R scripts for the DADA2 pipeline, subsequent analysis and visualization, along with the necessary data files are available at https://github.com/tpriest0/FRAM_EGC_2016_2020_data_analysis.

Co-occurrence networks were calculated for MIZ and core-EGC samples separately using the packages segmenTier^74^ and igraph^75^ in RStudio, and visualized in Cytoscape^76^ with the Edge-weighted Spring-Embedded Layout - detailed pipeline is available in Supplementary Information S1 and at https://github.com/tpriest0/FRAM_EGC_2016_2020_data_analysis. Briefly, oscillation signals were calculated for each ASV for each year based on discrete Fourier transformation of normalized abundances. Oscillation signals were then used to calculate the Pearson correlation between all ASVs, only retaining positive correlations. A network robustness analysis was performed to determine the minimal correlation value that represents a strong co-occurrence (0.7). Below this value, removal of a single node would cause network disruption. Full networks were built including only above-threshold co-occurrences. Values of centrality and node betweenness were calculated using igraph.

### PacBio metagenome sequencing

Nine samples from the 2016 – 2017 annual cycle at the MIZ were selected for metagenomic sequencing, using the same DNA as used for amplicon sequencing. Sequencing libraries were prepared following the protocol “Procedure & Checklist – Preparing HiFi SMRTbell® Libraries from Ultra-Low DNA Input” (PacBio, CA, USA) and subsequently inspected using a FEMTOpulse. Libraries were sequenced on 8M SMRT cells on a Sequell II platform for 30 h with sequencing chemistry 2.0 and binding kit 2.0. The sequencing was performed in conjunction with samples of another project, such that seven samples were multiplexed per SMRT cell. This resulted in, on average, 268 000 reads per metagenome, with an N50 of 6.8 kbp.

### Taxonomic and functional annotation of HiFi reads

The 2.4 million generated HiFi reads were processed through a custom taxonomic classification and functional annotation pipeline (https://github.com/tpriest0/FRAM_EGC_2016_2020_data_analysis). The classification pipeline followed similar steps to previously published tools, but with some modifications. A local database was constructed based on protein sequences from all species-representatives in the GTDB r202 database^77^. Prodigal v2.6.3^78^ was used to predict open reading frames (ORF) on HiFi reads, which were subsequently aligned to the GTDB-based database using Diamond blastp v2.0.14^79^ with the following parameters: --id 50 --query-cover 60 --top 5 --fast. After inspection of the hits, a second filtering step was performed: percentage identity of >65% and an e-value threshold of 1E-10. Using Taxonkit v0.10.1^80^, the last common ancestor (LCA) algorithm was performed, resulting in a single taxonomy for each ORF. A secondary LCA was subsequently performed for all ORFs from the same HiFi read, generating a single taxonomy for each read. Functional annotation of HiFi reads was performed using an extensive set of general and specialised databases. In brief, an initial gene annotation was performed using Prokka^81^. Then, a set of specialised databases were searched using blastp v2.11.0^82^ and HMMscan (HMMER v3.2.1)^83^ for further gene annotations, including dbCAN v10^84^, CAZy (release 09242021)^85^, SulfAtlas v1.3^86^, the Transporter Classification database^87^, MEROPS^88^, KEGG^89^ and sets of Pfam HMM family profiles for SusD and TonB-dependent transporter genes. In order to compare functional gene composition across samples, gene counts were normalised by the average count of 16 universal, single-copy ribosomal proteins per sample^90^ – providing ‘per genome’ counts.

### Metagenome-assembled genome recovery

In order to maximise the recovery of metagenome-assembled genomes (MAGs), we clustered the metagenomes into two groups. The groups were determined based on the dissimilarity in ASV composition of the corresponding samples, and largely reflected samples of high- and low-ice cover. The samples were then individually assembled using metaFlye v2.8.3 (parameters: --meta --pacbio-hifi –keep-haplotypes --hifi-error 0.01). Contigs with a length of <10 kbp were removed and the remaining contigs were renamed to reflect the sample of origin. The resulting contigs from each group were concatenated into a single file. Coverage information, necessary for binning, was acquired through read recruitment of raw reads from all metagenomes to the contigs using Minimap2 v2.1^91^, using the ‘map-hifi’ preset. Contigs were binned using Vamb v3.0.2^92^ in multisplit mode using three different sets of parameters (set1: -l 32 -n 512 512, set2: -l 24 -n 384 384 and set3: -l 40 -n 768 768, as suggested by the authors). Completeness and contamination estimates of bins were determined using CheckM v1.1.3^93^ and those with >50% completeness and <10% contamination were manually refined using the Anvi’o interactive interface (Anvi’o v7)^94^. MAGs from both groups were combined and dereplicated at 98% average nucleotide identity using dRep v3.2.2^95^ (parameters: -comp 50 -con 5 -nc 0.50 -pa 0.85 -sa 0.98), resulting in 47 population-representative MAGs. A phylogenetic tree was reconstructed using the representative MAGs from this study and those recently published from the Fram Strait by Priest et al.^22^ following a procedure outlined previously^90^. Briefly, 16 single-copy universal ribosomal proteins were identified in each MAG using HMMsearch against the individual Pfam HMM family profiles and aligned using Muscle v3.8.15^96^. The alignments were trimmed using TrimAI v1.4.1^97^, concatenated and provided as an input to FastTree v2.1.0^98^. The tree was visualised and annotated in iToL^99^.

### Classification, abundance and distribution of MAGs

A dual taxonomic classification of MAGs was performed using single-copy marker and 16S rRNA genes. Firstly, MAGs were assigned a taxonomy using the GTDBtk tool v1.7.0^100^ with the GTDB r202 database. Secondly, extracted 16S rRNA gene sequences were imported into ARB^101^, aligned with SINA^102^ and phylogenetically placed into the SILVA SSU 138 Ref NR99 reference tree using ARB parsimony. Those containing a 16S rRNA gene were linked to ASV sequences through competitive read recruitment using BBMap of the BBtools program v35.14, with an identity threshold of 100%.

The distribution of MAGs across the nine metagenome samples generated for this study and an additional 42 Arctic metagenomes from the Tara Oceans collection (Project: PRJEB9740) was determined through read recruitment. Counts of competitively mapped reads were converted into the 80% truncated average sequencing depth, TAD80^103^. Relative abundance was then determined as the quotient between the TAD80 and the average sequencing depth of 16 single copy ribosomal proteins. Ribosomal proteins were identified following the same procedure outlined above and their sequencing depth estimated using read recruitment with minimap2, for the metagenomes of this study, and BBMap, for Tara Oceans metagenomes.

### Mooring and satellite data

To place bacterial community data into context, we incorporated a collection of in situ environmental parameters that are presented in Supplementary Table S1. Temperature, depth, salinity and oxygen concentration were measured using Seabird SBE37-ODO CTD sensors and chlorophyll *a* concentration was measured using a WET Labs ECO Triplet sensor attached to the RAS. Sensor measurements were averaged over 4 h around each sampling event. These parameters were subsequently used to determine the relative proportions of Atlantic Water (AW) and Polar Water (PW) as described previously by Wietz *et al*.^23^. Physical sensors were manufacturer-calibrated and processed in accordance with https://epic.awi.de/id/eprint/43137. Mooring-derived data are published under Pangaea accession 904565^104^ and 941159^105^. Sea ice concentrations, derived from the AMSR-2 satellite, were downloaded from the https://seaice.uni-bremen.de/sea-ice-concentration-amsr-eamsr2, and averaged for the mooring regions using a 15 km radius.

## Data availability

The 16S amplicon sequences are available at EBI-ENA under the project accessions PRJEB43890 (2016-17), PRJEB43889 (2017-18), PRJEB54562 (2018-19), PRJEB54586 (2019-20). Individual sample accessions are provided in Supplementary Table S5. The metagenomic sequence data and MAGs generated for this study are available at EBI-ENA under the project PRJEB52171. Supplementary table S6 contains the respective accession numbers for the individual metagenomic raw read datasets, assemblies and MAGs used in this study. Functional gene annotations for all signature populations are provided in Supplementary Files S1. Physicochemical parameters used in this study are available under the Pangaea accession 904565^104^ and 941159^105^.

Code for reproducing analysis and generating figures, along with raw data files are available under https://github.com/tpriest0/FRAM_EGC_2016_2020_data_analysis.

## Acknowledgements

We thank Jana Bäger, Theresa Hargesheimer, Rafael Stiens and Lili Hufnagel for RAS operation; Daniel Scholz for RAS and sensor operations; Normen Lochthofen, Janine Ludszuweit, Lennard Frommhold and Jonas Hagemann for mooring operation; Jakob Barz, Swantje Rogge and Anja Nicolaus for DNA extraction and library preparation, and Bruno Huettel and the technicians at the Max Planck Genome Centre in Cologne for metagenome sequencing. The captain, crew and scientists of RV Polarstern cruises PS99.2, PS107, PS114, PS121 and PS126 are gratefully acknowledged. We thank Christina Bienhold, Katja Metfies, Oliver Ebenhöh and Eva-Maria Nöthig for helpful discussions. This project has received funding from the European Research Council (ERC) under the European Union’s Seventh Framework Program (FP7/2007-2013) research project ABYSS (Grant Agreement no. 294757) to AB. Additional funding came from the Helmholtz Association, specifically for the FRAM infrastructure, and from the Max Planck Society.

## Author contribution statement

TP performed ASV and metagenomics analysis. MW processed amplicon raw data into ASVs and coordinated the data analysis. TP and MW wrote the paper. WJvA contributed quality-controlled oceanographic data, and coordinated the mooring operations. EO and OP performed network analyses. STV provided quality-controlled chlorophyll sensor data. CB, KM and AB co-designed and coordinated the autonomous sampling strategy, and contributed to interpretation of results. BF and RA contributed to interpretation of results and development of the story. All authors contributed to the final manuscript.

