## Supplementary figures and images for "Variations in Atlantic water influx and sea-ice cover drive taxonomic and functional shifts in Arctic marine bacterial communities"

### Supplementary Figure S1

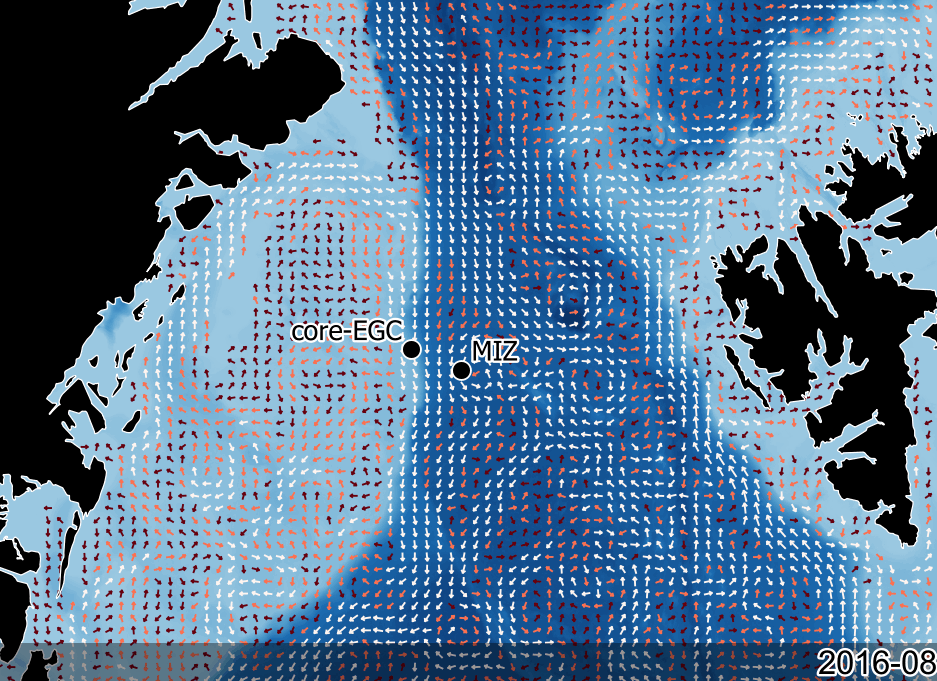

### Supplementary Figure S2

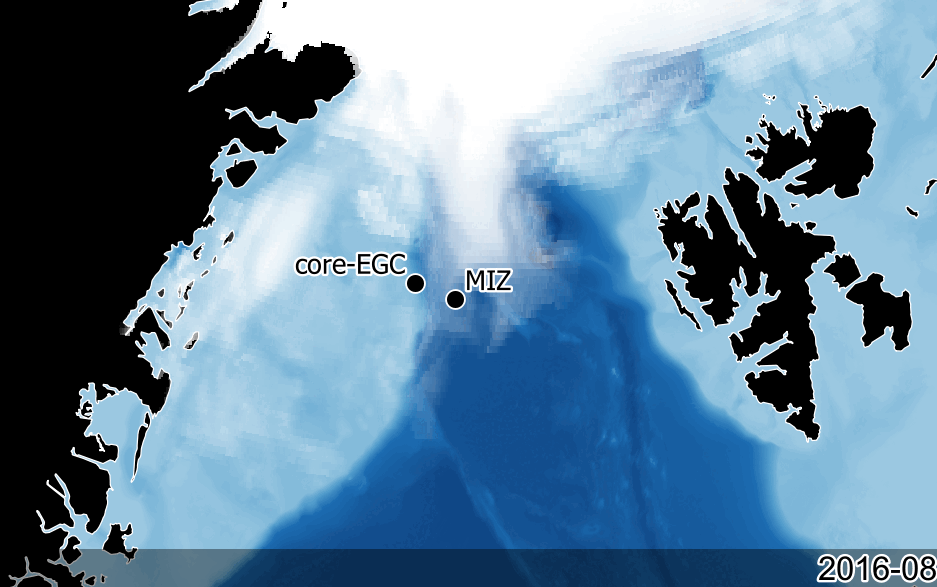

### Supplementary Figure S3

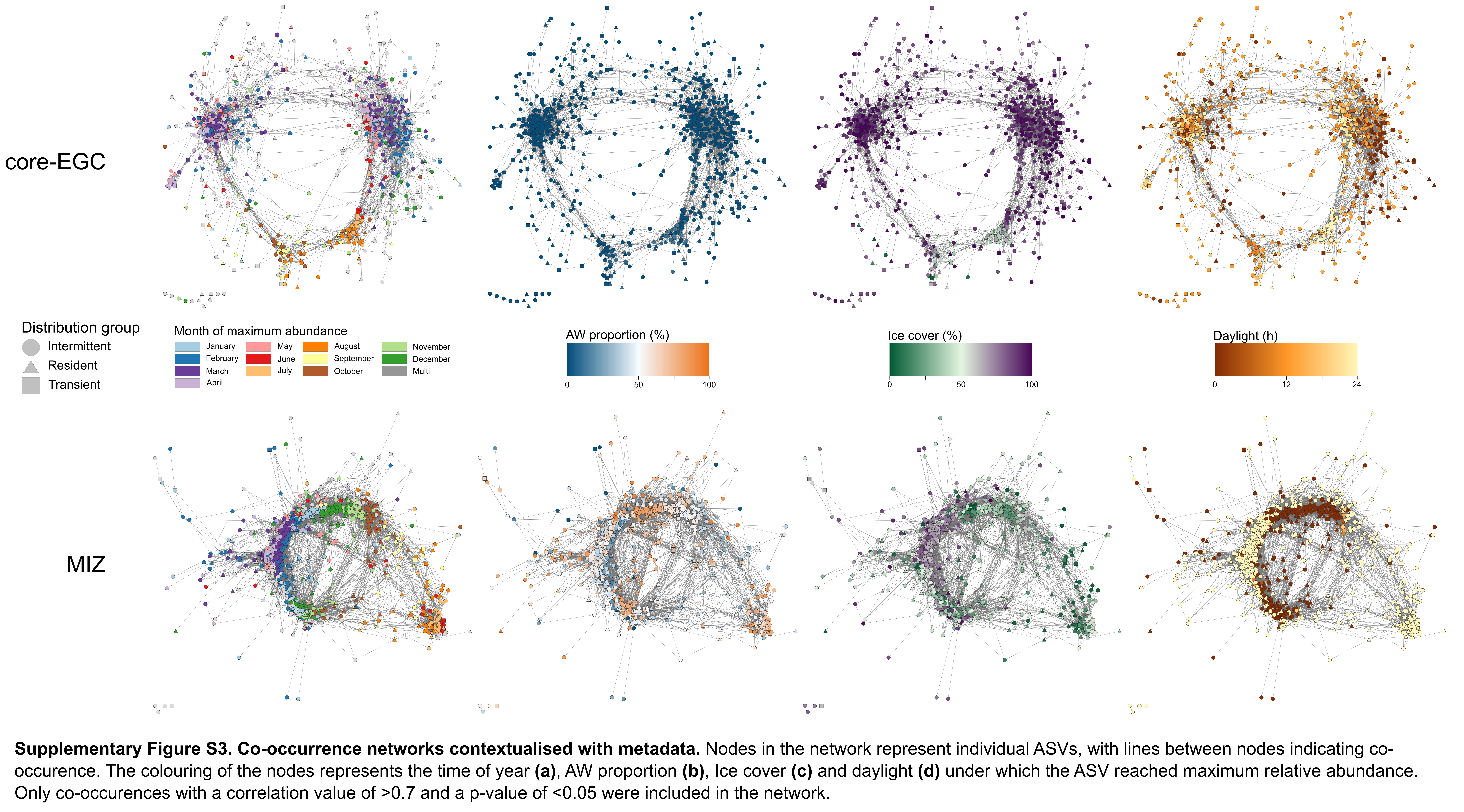

### Supplementary Figure S4

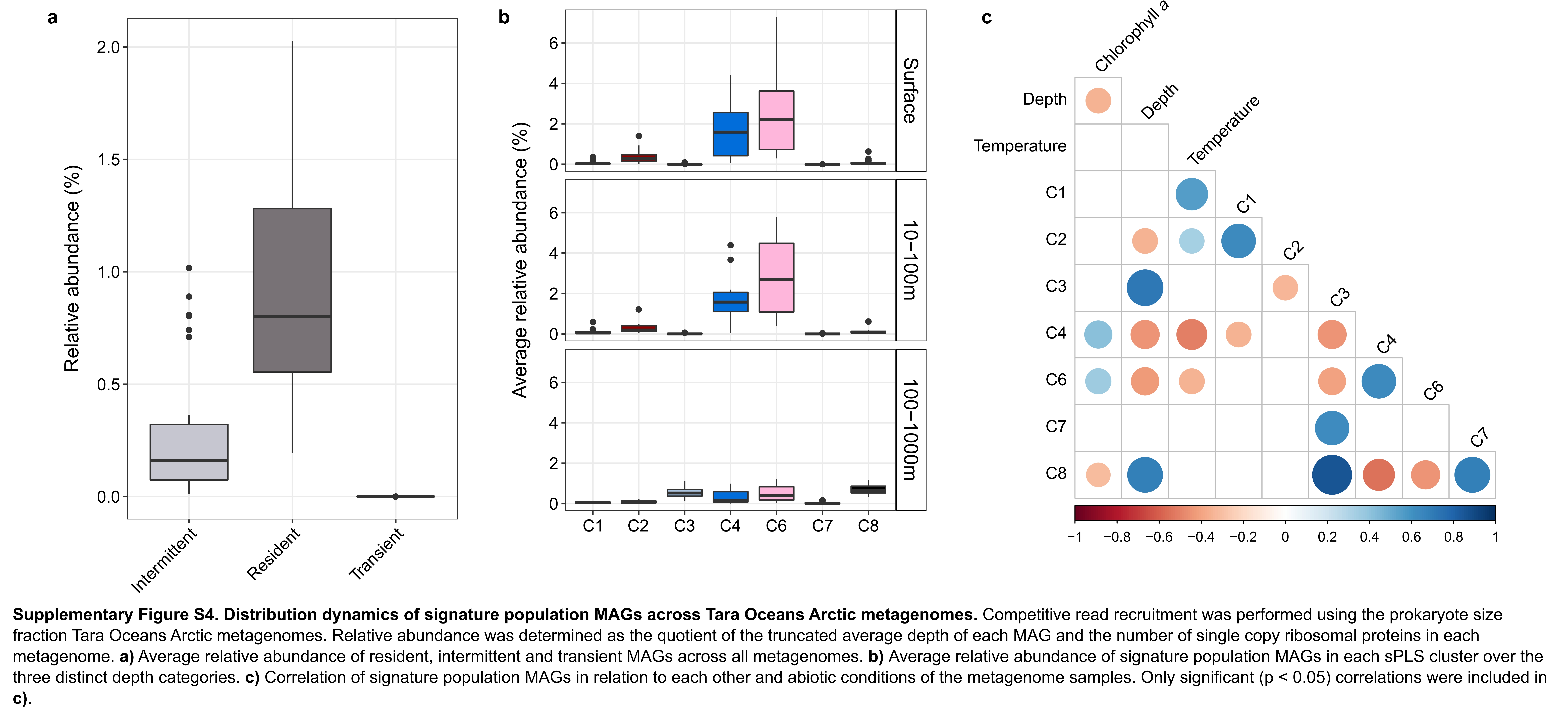

### Supplementary Figure S5

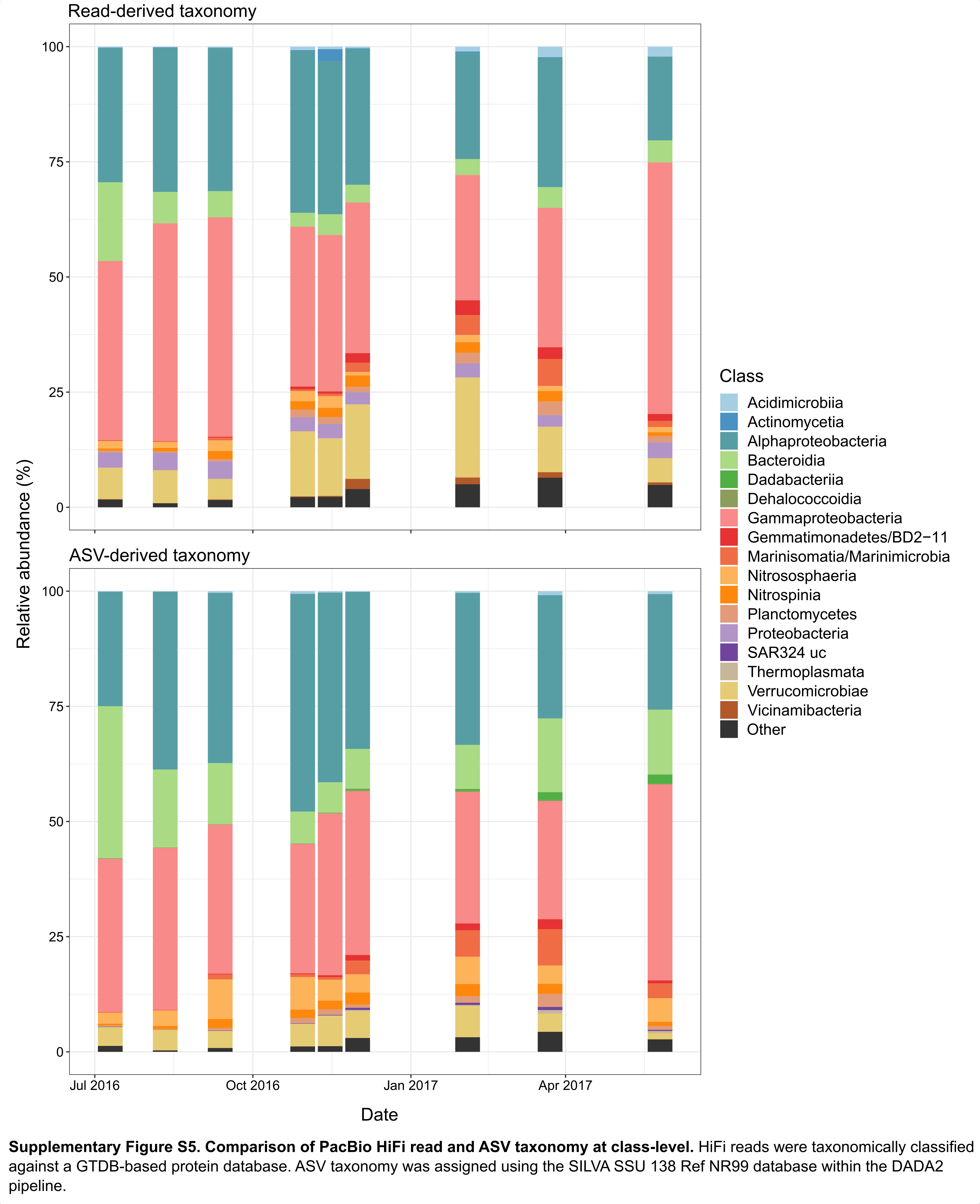

### Supplementary Figure S6

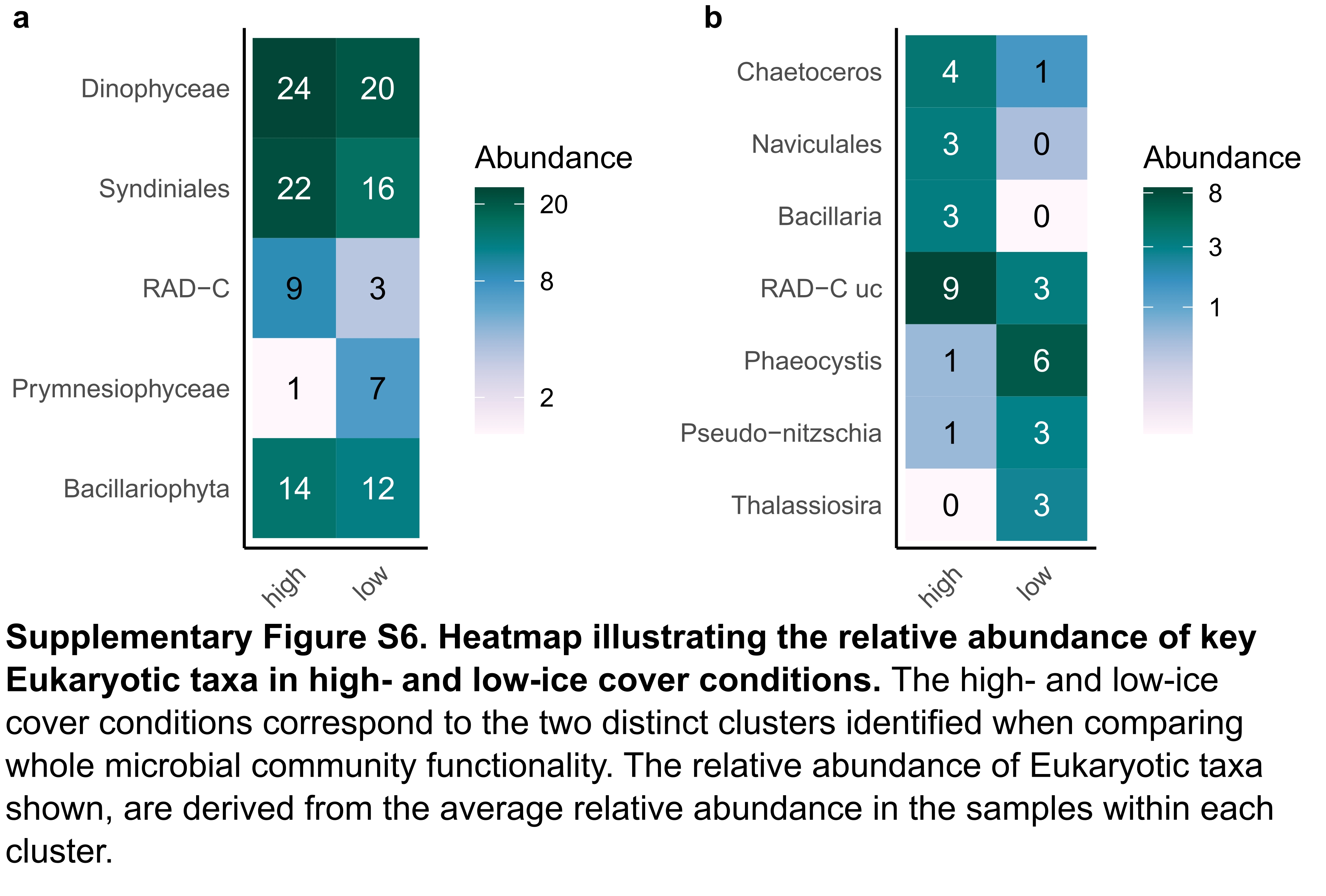
