## Supplementary Information S1 for "Variations in Atlantic water influx and sea-ice cover drive taxonomic and functional shifts in Arctic marine bacterial communities"

**SUPPLEMENTARY RESULTS**

*Ecological niches of signature populations*

By combining temporal dynamics with functional gene predictions, we are able to make predictions on the ecological niches of signature populations within the context of the environmental conditions they represent. Signature populations of interest from clusters C1, C2 and C5 were characterized in detail in the main manuscript text, however additional signature populations from cluster C5 and those from C3, C4 and C6 were not described. Here, we provide ecological descriptions for the remaining signature populations.

*Remaining Arctic signature populations (cluster C5)*

The remaining Arctic signature populations were affiliated with *Nitrospina* (asv118) and the OM75 Clade (asv163), both of which exhibited similar dynamics and represented comparable proportions of the communities, reaching 1.7 – 1.8% relative abundance values. These populations harboured distinct functional features, relying on different substrates for growth but also performing key processes that contribute to the cycling of carbon, nitrogen and sulfur under high-ice coverage conditions.

The asv163 population was assigned to the OM75 Clade in the SILVA database and the GCA-2722775 (within the Nisaeaceae) in the GTDB database. Functional gene annotations indicated a non-motile lifestyle with a photoheterotrophic metabolism that included a green-light proteorhodopsin and the capacity to use a diverse repertoire of organic substrates for C, N and S acquisition and energy generation. Of particular note, was the extensive set of ABC transporters, 53, compared to other Arctic signature populations, which contained, on average, 17. Exogenous carbon substrates likely include osmolytes, C1 and aromatic compounds. The asv163 population encoded the C1 tetrahydrofolate oxidation pathway, similar to some members of the SAR11 clade ^1^, which also produces energy through ATP. The degradative pathways for the osmolytes taurine, sarcosine and choline were encoded, whilst partial metabolic pathways for aromatic compound metabolism were also identified. Organic compounds also act as the main sulfur source for the asv163 population, with the degradation of sulfonates through 2-aminoacetaldehyde to sulfite. The sulfite could subsequently be converted to sulfate through the sulfite dehydrogenase gene (*soeABC*) and assimilated through 3’-phosphoadenylyl-sulfate. In addition to sulfonates, the asv163 population also harboured the capacity to utilise dimethylsulfide, dimethylsulfoxide and dimethylsulfone, which can act as a sulfur and carbon source, with the production of formaldehyde being channeled through the tetrahydrofolate oxidation pathway. Organic nitrogen sources for the asv163 population are predicted to include amino acids and urea. Several branched-chain and L-amino acid transporters were unique to this population as well as the complete urease enzyme complex. The asv163 population thus harbours an extensive capacity to uptake and degrade diverse organic substrates, which would be advantageous under high-ice conditions where the input of fresh organic matter is limited.

The asv118 population, assigned to the *Nitrospina* genus in SILVA and SCGCAAA288-L16 in GTDB, could be described as motile, chemolithotrophic organisms, in line with descriptions from previous studies ^2,3^. *Nitrospina* belong to the *Nitrospinota* phylum that comprises the most abundant nitrite oxidising bacteria (NOB) in the oceans. Identified from surface to deep waters and from oxygenated to oxygen minimum zones, *Nitrospinota* are essential for the marine nitrogen cycle, with microbial nitrite oxidation reported to be the most significant biological pathway for nitrate production in the oceans^4^. However, the essential nitrite reductase, *nirK* gene, was not identified in the asv118 population, which may reflect the lower completeness of the MAG. The presence of a nitrite:ferredoxin reductase, *nirA*, indicates a capacity to convert nitrite to ammonia in an assimilation process and reflects previous obvious in other nitrite oxidizing bacteria of the *Nitrospira* genera^5^. Additional routes for nitrogen acquisition included ammonium uptake and a capacity to use urea through the *ureABC* gene. Additional observations in the asv118 population MAG included a green-light proteorhodopsin, suggesting supplemental energy generation through light - a feature not previously reported for *Nitrospina* members. Although the key nitrite oxidation machinery was lacking, we hypothesise that this process would still formulate a key part of energy generation in this organism, due to its highly conserved nature across NOB taxa. Recently, evidence for hydrogen oxidation in NOB taxa was reported, however no such genes were identified in the asv118 representative MAG.

*Polar day signature populations (clusters C3 and C6)*

Seven signature populations were identified for polar day conditions, six assigned to the Atlantic water-associated cluster C3 and one to the Arctic water-associated C6. Despite the affiliations to different sPLS clusters, C3 signature populations also reached comparable relative abundance values at the core-EGC under polar day conditions, suggesting that water mass is less influential. In contrast, the asv16 (SAR92 Clade) Arctic-water associated population did not reach comparably high relative abundances in MIZ samples, suggesting that this population is representative of polar day conditions only in Arctic water masses. Furthermore, the dynamics of the populations across the three polar day time periods in the core-EGC mooring appeared to be dependent on the magnitude of chlorophyll *a* concentrations measured; this pattern indicates an intrinsic link to phytoplankton dynamics and suggests that a certain threshold of phytoplankton abundance must be reached before such a response is observed in these populations. Each of the polar day signature populations identified here are affiliated with taxa that are well-known to be responders to phytoplankton blooms in marine environments ^6,7^. At Helgoland Roads, a swift and recurrent succession of bacterial clades following phytoplankton blooms has been observed, with consecutive peaks of *Ulvibacter*, *Formosa, Reinekea, Polaribacter* and SAR92 from the *Bacteroidia* and *Gammaproteobacteria* class^8^. Here, the ephemeral peaks in relative abundance of some of these clades exhibited a more coordinated and less successional pattern, however this may reflect the lower resolution of sampling. In general, asv6 (*Polaribacter*), asv7 (*Aurantivirga*), asv55 (*Ulvibacter*), asv71/88 (*Nitrincolaceae*) and asv16 (SAR92) all exhibited rapid increases in relative abundance, peaking at the same time point followed by a decrease over a 4 week period. Although no chlorophyll *a* data is available from the MIZ mooring, the dynamics observed at core-EGC would indicate that these patterns are a response to phytoplankton blooms. As such, we hypothesise that these populations are involved in the degradation of phytoplankton-derived organic matter, but each occupying distinct substrate-based niches, as has been observed at Helgoland Roads.

*Aurantivirga* and *Polaribacter* have been shown to harbour broad substrate utilisation capacities but also occupy distinct polysaccharide-based niches^9,10^. In accordance with previous findings, both the asv6 (*Polaribacter*) and asv7 (*Aurantivirga*) representative MAGs harboured rich CAZyme gene repertoires and polysaccharide utilisation loci (PULs) for carbohydrate degradation. This consisted of 31 and 28 degradative CAZymes in asv6 and asv7, respectively, along with three distinct PULs in each. Two PULs identified in both of the populations are predicted to target the diatom storage polysaccharide laminarin (PUL 1 containing GH16_3, GH17 and GH149 and PUL 2 containing GH16_3 and GH30_1). The third PUL in the asv7 population is predicted to target α-glucans (GH13, GH13_31 and maltose transporter) whilst in asv6, sulphated xylose-containing polysaccharides are the predicted target for the third PUL (GH10, GH113, several sulfatases and D-xylose transporter). The gene synteny and structures of the PULs are also in agreement with those previously described for *Polaribacter* and *Aurantivirga* representatives from Helgoland Roads^9^. Further comparisons revealed species-level differences in the dominant populations identified in our dataset and that of Helgoland Roads. For example, the asv6 Polaribacter population MAG shares 94.9% amino acid identity with 20120426_Bin_74_1 from Helgoland Roads (PRJEB28156), which was described as being present only in particular seasons and years but not related to the *Polaribacter* species that dominates during spring phytoplankton blooms. The above-described metabolic capacity, combined with ephemeral but pronounced peaks under polar day and alongside chlorophyll *a* peaks in the EGC, indicate the occupation of distinct substrate-based ecological niches for the asv6 and as7 populations. Furthermore, the comparable relative abundance values reached at both core-EGC and MIZ suggest that these populations are capable of proliferating under contrasting conditions, suggesting that substrate availability is the key factor defining their ecological niche.

In addition to the coordinated dynamics observed for members of the *Bacteroidia* and *Gammaproteobacteria*, three signature populations affiliated with the *Verrucomicrobiae* also exhibited pronounced increases in relative abundance. These populations were affiliated with the BACL24 (*Lentimonas*) and UBA1315 (*Luteolibacter*) genera. However, the dynamics of these populations differed. The asv94 population (*Lentimonas*) peaked with the *Bacteroidia* and *Gammaproteobacteria* representatives at core-EGC but showed a more delayed response at MIZ whilst the asv24 and asv115 (*Luteolibacter*) populations typically peaked later. These variations likely reflect the occupation of different ecological niches. Members of the *Verrucomicrobiae* are well evidenced to respond to phytoplankton blooms and are typically described as degraders of more complex polysaccharide structures, particularly those that are heavily sulfated^11,12^. In accordance with previous findings, the *Luteolibacter* representatives encoded a large number of degradative CAZymes, 40 in asv115 and 31 in asv24, and a high sulfatase to CAZyme ratio, 1:0.8 in asv115 and 1:0.7 in asv24. Further analysis on the encoded CAZyme genes revealed key distinctions between these two populations. Asv24 encoded genes assigned to five CAZyme gene families that are known to target alpha-glucans/amylose (GH13_38, GH13_4, GH13_8, GH57 and GH77) compared to only one gene in the asv115 (GH13_38). In addition, the asv115 population encoded two CAZymes that target rhamnogalacturonan (GH105 and GH106) which were absent from asv24. These metabolic distinctions may contribute to explaining the large difference in maximum relative abundances observed between these populations (6.1% in asv115 and 15.3% in asv24) and point towards substrate-based niche partitioning. Interestingly, the *Verrucomicrobiae* that are known to be the most prominent responders to spring phytoplankton blooms at Helgoland Roads are not from the *Luteolibacter* genera, but affiliated with different genera of the *Akkermansiaceae* family or with the *Lentimonas* genus of the *Puniceicoccaceae* family^12^. In contrast, the *Lentimonas* population here (asv94) reached much lower relative abundances than the *Luteolibacter* populations. This further illustrates differences in the microbial populations that respond to phytoplankton blooms in different ecosystems.

In difference to the above-described polar day-associated representatives, the asv71/88 population harboured distinct metabolic features, including a capacity for methylotrophy and a rich genetic repertoire for sulfur metabolism. Classified as *Nitrincolaceae* in the SILVA database and assigned to the ASP10-02a in GTDB, we describe the asv71/88 population as a motile chemoheterotroph. Methylotrophic metabolism was evidenced by genes involved in trimethylamine utilisation (*tmm, dmd-tmd, mgsABC* and *mbdAB*), with the produced formaldehyde likely being converting to CO2 through formate (*fdoG* and *fdwB*) and the ammonium being used as a nitrogen source. The asv71/88 population encoded a large number of genes involved in sulfur metabolism that included the ability to use organic sulfur compounds (methanesulfonate and sulfopyruvate) along with the complete thiosulfate oxidation machinery (*soxABCDXYZ*) and a sulfite dehydrogenase (*soeABC*). Methanesulfonate (MSA) is one of the main products of dimethylsulfide oxidation, and thus is likely present at higher concentrations during polar day conditions and phytoplankton blooms. The oxidation of MSA through a MSA monooxygenase, encoded in asv71/88 population, results in the production of formaldehyde and sulfite, which can be further oxidised through energy-generating reactions to CO2 and sulfate. Alongside the capacity to use organic nitrogen and sulfur compounds, the asv71/88 population also harboured 26 ABC transporters, including those for amino acid, monosaccharides, polyols and urea uptake, as well as the potential to degrade toluene. Furthermore, we identified the genes for a mannose-sensitive haemagglutinin-like pilus (*mshCDGIJKLOP*), which has been shown to promote attachment of bacterial cells to algae^13^. Therefore, the ability for motility and attachment combined with a diverse metabolic capacity of the asv71/88 population indicates a copiotrophic lifestyle that could involve a close relationship with phytoplankton cells, similar to that described for *Vibrio* and *Pseudoalteromonas* representatives. The encoded pathways for biotin, riboflavin, cobalamin and pantothenate synthesis, suggest that vitamins may be a valuable product provided to the phytoplankton by the asv71/88 population.

*Polar night signature populations (clusters C4)*

Signature populations of polar night conditions consisted of asv13, assigned to the *Arenicellaceae* family in SILVA and UBA11654 in GTDB, and asv8, assigned to the Arctic97B-4 in SILVA and UBA1096 in GTDB. The dynamics of these two populations were largely consistent, with highest relative abundance values of 4.6% for asv13 and 3.8% for asv8 reached under polar night conditions in MIZ samples. Insights into the metabolic capacity and potential ecological niches of these populations revealed some distinctions, however the asv13 representative MAG was of lower completeness, 63%, which was reflected in the annotation of incomplete pathways that hindered the analysis.

Arenicellaceae is a family of Gammaproteobacteria that has previously been reported in deep-sea sediments^14^, responding to phytoplankton blooms in coastal seawater^15^ and has been proposed as an indicator of eutrophication^16^. The limited ecological information on members of this family indicates an organotrophic lifestyle. Due to the lower completeness of this MAG, we cannot provide clear predictions on the ecological niche but will briefly outline key metabolic features found. In general, the asv13 population could be characterized as non-motile and harbouring a capacity to use C1 compounds as substrates for growth along with indications of nitrate reduction (nitrate reductase, *narH*) and carbon fixation (incomplete rTCA cycle). C1 metabolism was indicated by the presence of a methanol dehydrogenase, formate dehydrogenase, methylenetetrahydrofolate dehydrogenase (*folD*) and the complete pathway for cofactor F420 biosynthesis. The presence of a nitrate reductase, *narH*, suggests a potential for nitrate reduction, however the other key subunits were missing. Two annotated ammonium channels also highlighted additional routes for nitrogen acquisition. We further focused on transport systems to reveal additional information on substrates used for growth, but those identified included typical transporters that are widespread in marine bacteria, such as vitamin B12, magnesium, sialic acid and general biopolymer transport (*exbBD*). Furthermore, we identified a complete riboflavin biosynthesis pathway. As a result, the ecological role of the asv13 population under polar night conditions is yet unknown and further analysis is required.

The asv8 population was assigned to the same taxonomic group in SILVA, Arctic97B-4, as the Arctic signature population (asv191), however the GTDB classification places these two populations into distinct families. This phylogenetic difference is also supported by distinctions in lifestyle and metabolic capacities. Indifference to the Arctic signature population, the asv8 population can be categorised as non-motile and harbouring an extensive genetic repertoire for organic compound metabolism, particularly carbohydrates. A total of 55 degradative CAZymes and 54 sulfatases were identified, highlighting a rich carbohydrate degradation capacity, which includes substrates such as pectin/rhamnogalacturonan (PL1 x 2, GH165 x 2, GH140 and CBM67 x 2), β-glucuronyl-containing polysaccharides (GH88, CE15 x 2) and sialic acids (GH33 x 3). Also encoded within the asv8 population was the capacity to use additional organic substrates as carbon and energy sources, which included glyoxylate and dicarboxylates, such as glycolate (*glcDEF*). For the acquisition of nitrogen and sulfur, inorganic sources are likely also used, with the complete sulfate assimilation pathway present and a nitrate reductase gene (*napA*) – although in the case of nitrate reduction not all necessary subunits and genes were present. Another key metabolic feature, which likely proves advantageous during winter conditions when fresh organic substrates are scarcely available, was the ability to synthesise glycogen (GBE1 and *glgACE*). Glycogen is a sugar compound that plays important roles in energy and carbon storage in some bacteria and has been shown to increase bacterial durability under starvation conditions^17^. Although the internal hydrolysis of glycogen would not provide sufficient resources to explain observed dynamics during winter, it could certainly aid in population preservation under unfavourable conditions. With this, it could be hypothesized that the asv8 population is able to survive through the use of inorganic substrates and complex, recalcitrant carbohydrate substrates in winter conditions. The complex carbohydrates may be residual compounds left over from polar day conditions but may also be derived from under-ice algae and/or terrestrial-derived DOM.
